# Decoding efficacy and resistance space at a drug binding site

**DOI:** 10.1101/2025.07.25.666894

**Authors:** Simone Altmann, Cesar Mendoza-Martinez, Melanie Ridgway, Michele Tinti, Jagmohan S. Saini, Peter E. G. F. Ibrahim, Michael Thomas, Manu De Rycker, Michael J. Bodkin, David Horn

## Abstract

Interactions between drugs and their targets impact efficacy and, when altered by mutation, can result in resistance ^1-3^. Assessing and understanding the impacts of all possible mutations at a drug binding site remain challenging, however ^4-6^. Here, we used Multiplex Oligo Targeting (MOT) for mutational profiling, and computational modelling, to decode efficacy and resistance space at the otherwise native binding site for a low nanomolar potency, anti-trypanosomal, proteasome inhibitor ^7^. We saturation-edited twenty codons in the *Trypanosoma brucei* proteasome β5 subunit and subjected the resulting MOT libraries to stepwise drug selection. Amplicon sequencing, and codon variant scoring, yielding dose-response profiles for >100 resistance-conferring mutants, among 1,280 possible codon variants. Codon variant scores were predictive of relative resistance observed using a bespoke set of mutants, while fitness profiling revealed otherwise extensive constraints on mutational fitness and resistance space. The resistance profile that emerged allowed us to readily predict routes to spontaneous drug resistance observed within ‘accessible’, single nucleotide mutational space. *In silico* analysis of β5 subunit mutations predicted impacts on ligand affinity via steric effects, hydrogen-bonding and lipophilicity, which when combined with predictions of proteasome function perturbing mutations, were closely aligned with observed impacts on drug resistance. We conclude that MOT-library profiling facilitates assessment of all possible mutations at a drug binding site. Further decoding of drug target structure-activity relationships and drug resistance space will facilitate the design of more effective and durable drugs.

## Introduction

Improved understanding of structure-activity relationships between drugs and their targets facilitates progress in drug discovery. The emergence of drug resistance, on the other hand, undermines the durability of chemotherapeutics against infectious diseases and in oncology. Insights into structure-activity relationships typically emerge from a combination of empirical experimental observation *in cellulo,* structural studies, and computational modelling and prediction *in silico*. Notably, although powerful approaches for saturation mutagenesis, combined with multiplexed assays, are available, *in cellulo* data describing comprehensive sets of mutations at native drug binding sites are often lacking ^8^, and confidence in *in silico* prediction can suffer from unquantified limitations as a result.

Several CRISPR-based editing approaches have been used to assess drug resistance associated mutations in eukaryotic cells. However, these approaches employ Cas9 nucleases that can yield undesirable and imprecise recombinants, base editors that typically introduce <50 % of all possible edits ^5,9^, or prime editors that display limited efficiency ^4^. CRISPR-based approaches are also typically constrained by the availability of suitable protospacer-adjacent motifs in the editing window and necessitate the introduction of further changes at the target site to prevent multiple rounds of editing. These approaches also present multiplexing challenges in diploid cells since different edits will likely be introduced in each allele, complicating genotype to phenotype assessments; this latter challenge can be mitigated by using haploid cells ^10^. Deep mutational scans, widely used for multiplex analysis of variant effects ^11^, have also been used to assess drug resistance mechanisms. However, these approaches typically employ ectopic expression systems that may alter target protein abundance and/or expression patterns ^12^.

Alternative approaches for the delivery of codon edits to native genes are available. Indeed, editing using multiplex genome engineering ^13^, or oligo targeting ^14^, simply requires the delivery of short single-stranded oligonucleotides. These approaches are DNA cleavage-free, and CRISPR-independent, and can deliver the full range of possible base edits, albeit at relatively low frequency. These approaches have been described in *Saccharomyces cerevisiae* ^13^, in embryonic stem cells ^15^, and in the trypanosomatids, *Trypanosoma brucei, T. cruzi* and *Leishmania* ^14^; parasitic protozoa that are transmitted by insects, causing neglected tropical diseases in humans, and also a range of veterinary diseases.

The proteasome is a promising drug target in the trypanosomatids ^7,16–19^, in malaria parasites ^20^, and against multiple myeloma ^21,22^. Cryo-electron microscopy structures of the *Leishmania* proteasome reveal a common binding site for both GSK3494245 ^7^ and LXE408 ^23^, and anti-trypanosomal proteasome inhibitors are currently in clinical development against both visceral leishmaniasis, and *T. cruzi*, which causes Chagas disease. We recently reported the development of oligo targeting for the assessment of drug resistance mechanisms in the trypanosomatids ^14^, and have now scaled the approach for saturation-editing of the codons encoding a drug binding site in the *T. brucei* proteasome β5 subunit. Multiplex oligo targeting and codon variant scoring, combined with computational modelling, provides unprecedented insights into structure-activity relationships between a drug and a mutated, but otherwise native, drug target. The readouts reveal how each binding site residue, and all possible binding site mutations, impact drug efficacy and drug resistance.

## Results

### Saturation mutagenesis at the proteasome β5 drug binding site

The proteasome is a high-priority drug target and was selected for the development and testing of multiplexed oligo targeting in trypanosomes. The proteasome comprises four stacked heteroheptameric rings with the β subunits contributing six proteolytic active sites that reside in the central chamber. These sites are autocatalytically activated in the assembled structure via cleavage of N-terminal propeptides; at the glycine preceding the catalytic threonine in the *T. brucei* β5 subunit (Tb927.10.6080) ^24^. Anti-trypanosomal proteasome inhibitors selectively target the chymotrypsin-like site in this subunit ^7,16,23^.

Oligo targeting in *T. brucei* involves delivery of single-stranded oligodeoxynucleotides (ssODNs) of approx. 50 b in length, with higher efficiency editing achieved when using ‘reverse-strand’ ssODNs ^14^; a single allele is typically edited in *T. brucei*, which is diploid. To target the chymotrypsin-like site in the β5 subunit, we considered a pooled ssODNs delivery protocol, but noted the potential for introduction of multiple edits in the same cell, which would allow for enrichment of bystander edits, complicating codon-based genotype to phenotype assessments. We, therefore, tested co-editing efficiency by delivering pairs of ssODNs to wild-type *T. brucei* cells, one designed to introduce a resistance-conferring edit (G^98^S - *G*G*A*-*A*G*C*), equivalent to G^98^S in *T. cruzi* ^14^, and a second designed to introduce a synonymous edit either approx. 100 (C^63^C, TG*C*-TG*T*) or 200 bp (V^31^V, GT*T*-GT*A*) upstream. Although an allele replacement frequency of only 0.003 % was reported previously in wild-type *T. brucei* ^14^, not readily detectable using Sanger sequencing, we readily detected synonymous edits in drug-resistant populations generated using this approach (Supplementary Fig. 1), indicating efficient co-editing at adjacent sites. Accordingly, and to avoid co-editing, we subsequently delivered each degenerate ssODN individually.

We selected compound DDD247, a nanomolar potency inhibitor (compound 7a in ^7^), for detailed analysis, and used cryo-EM structural data ^7^ to identify amino acids located within 5 Å of the proteasome β5 subunit binding site (Fig. 1a). The cognate codons for these twenty amino acids were distributed over a region of approx. 500 bp in the *β5* gene. To assemble a DDD247 binding site mutant library, we designed twenty reverse-strand, 53-b, ssODNs, each with a centrally located degenerate ‘NNN’ (N=A, C, T, G) codon (Fig. 1b). Mismatch-repair, previously reported to supress editing efficiency approx. 50-fold ^14^, was transiently knocked down using RNA interference for 24 h and, to avoid co-editing at adjacent sites in individual cells (see above), each degenerate ssODN was delivered individually. We then pooled the resulting populations and split the pool to generate a pair of Multiplex Oligo Targeting (MOT) libraries. Given a calculated allele replacement frequency of >0.04 % using this approach (Supplementary Fig. 2), we estimated an average yield of >1,000 codon edits per ssODN. To assess editing at each of the targeted codons, we extracted genomic DNA before, and six hours after library assembly, PCR-amplified the edited region in the proteasome *β5* gene, and deep-sequenced the amplicons (Fig. 1b). A scan to quantify variant codons across the edited region revealed highly specific editing at all twenty targeted sites (Fig. 1c). We concluded that all 1,280 possible variants around the β5 subunit drug binding site were likely represented in our pooled MOT libraries. These results served to validate the edited MOT libraries and the codon variant scoring approach.

**Fig. 1:**
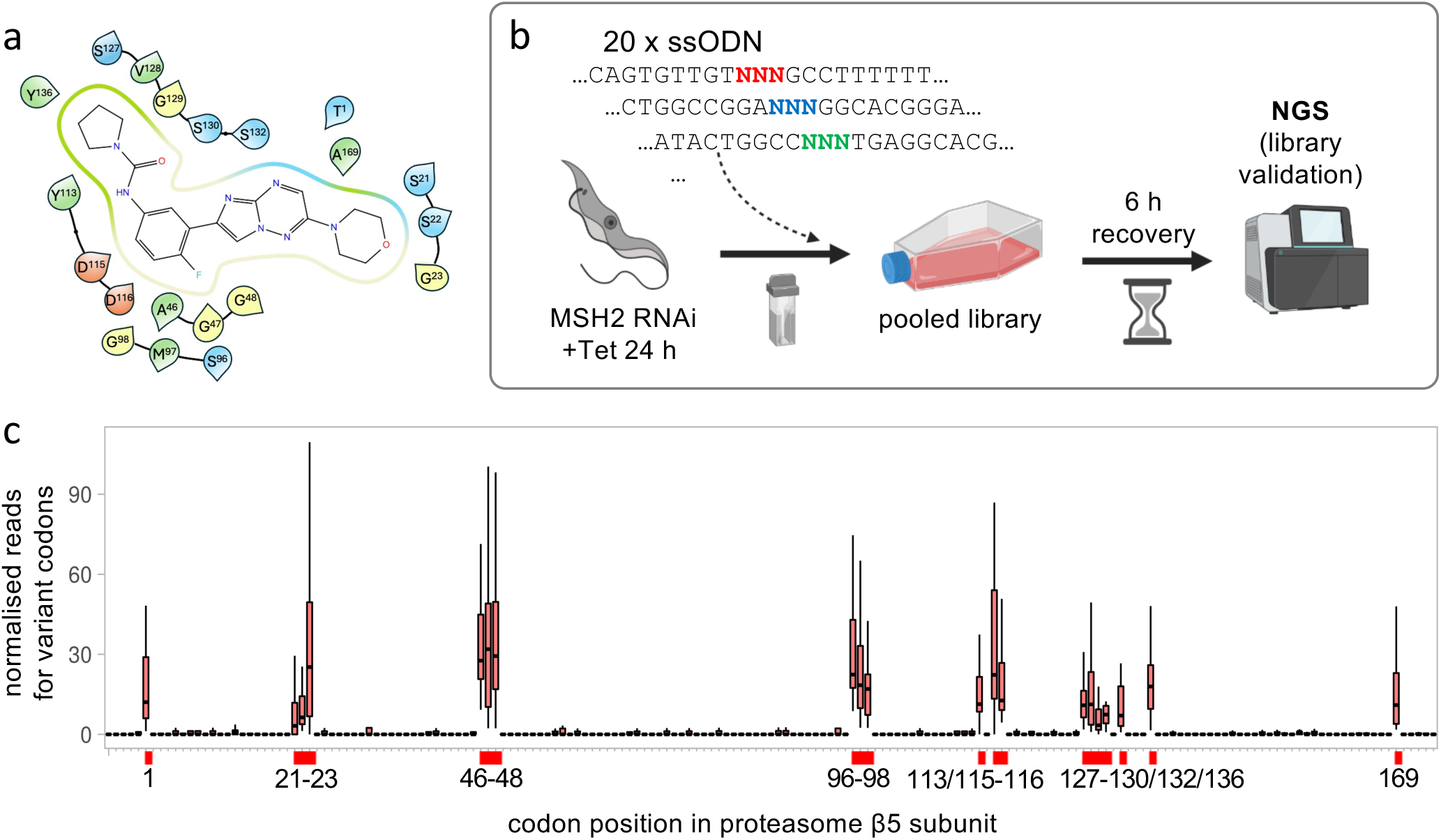
Saturation mutagenesis at the proteasome β5 drug binding site. **a** The ligand interaction diagram shows twenty *T. brucei* proteasome β5 subunit residues that are within 5Å of the docked compound, DDD247. Green, hydrophobic; blue, positively charged; red, negatively charged; yellow, glycine. **b** The schematic illustrates the MOT-library assembly approach. Twenty single-stranded oligodeoxynucleotides (ssODN) with a central degenerate codon were individually transfected into *T. brucei MSH2* RNAi cells 24 h after inducing knockdown with tetracycline. After 6 h, the pooled culture was assessed using deep sequencing of β5 amplicons. Created in BioRender. Altmann, S. (2025) https://BioRender.com/3zjhvug **c** The boxplot shows specific editing for all twenty targeted codons, indicated on the x-axis, and as determined by deep sequencing and codon variant scoring. Boxes indicate the interquartile range (IQR), the whiskers show the range of values within 1.5 × IQR, and a horizontal line indicates the median. Codon numbering is based on the mature β5 peptide.

### Codon variant scoring reveals >100 resistance-conferring base edits

To select drug-resistant mutants, the MOT libraries were grown in compound DDD247 at 8 nM; twice the EC_50_ (Effective Concentration of drug to inhibit growth by 50 %). The selected libraries were then split three ways and grown in compound DDD247 at either 40, 200 or 1,000 nM (Fig. 2a). We extracted genomic DNA following selection at each drug concentration, PCR-amplified the edited region in the proteasome *β5* gene, deep-sequenced the products (Fig. 2a), and quantified variant codons. Principle Component Analysis of variant codon read counts revealed similar profiles for the replicate MOT libraries (Fig. 2b). An assessment of edits generating stop/nonsense codons (n = 60) or synonymous codons (n = 61), neither of which were expected to yield resistant cells, revealed negligible read counts across the dose-response profile (Fig. 2c). In contrast, a number of edits generating mis-sense (non-synonymous) codons yielded elevated read counts across the dose-response profile, at the Y^113^ codon for example (Fig. 2c).

**Fig. 2:**
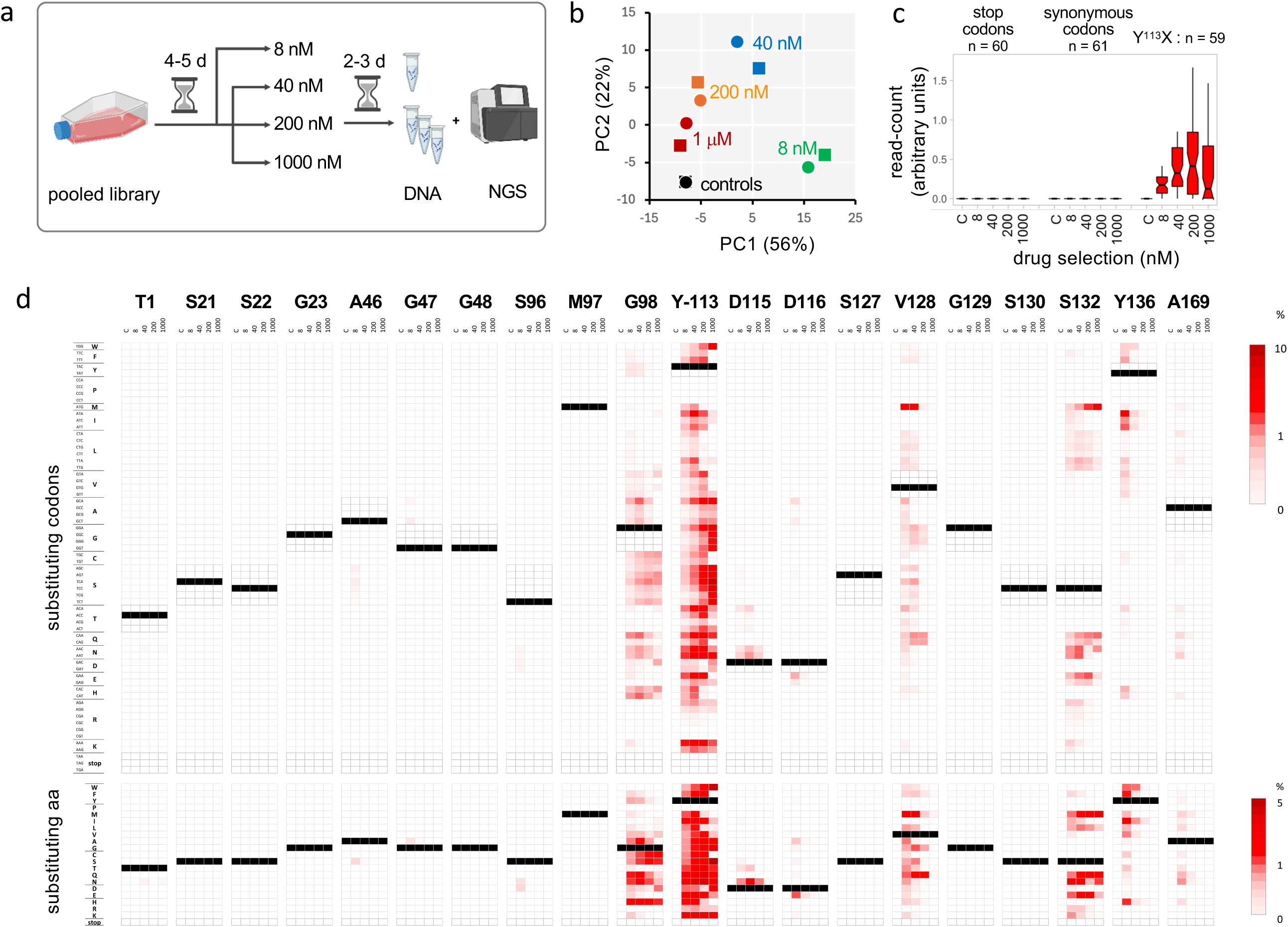
Codon variant scoring reveals >100 resistance-conferring base edits. **a** The schematic illustrates the MOT-library profiling approach. The edited, pooled *T. brucei* library was split in duplicates, which were subjected to compound DDD247 selection as indicated. Genomic DNA was then isolated, and *β5* gene amplicons were analysed by deep sequencing and codon variant scoring. Created in BioRender. Altmann, S. (2025) https://BioRender.com/zy1spe1 **b** Principle Component Analysis indicated that codon variant scores diverged from the unselected control during DDD247-selection and that the replicate experiments yielded consistent results. **c** The boxplot shows codon variant scores for the indicated categories and across the dose-response profile. Boxes indicate the interquartile range (IQR), the whiskers show the range of values within 1.5 × IQR, and a horizontal line indicates the median. The notches represent the 95% confidence interval for each median. **d.** All sixty-four possible codon variant scores for all twenty targeted sites in the proteasome *β5* subunit gene are represented as a heatmap; for the unselected control sample, and across the dose-response profile (upper panel). A heatmap for amino acid variant scores is shown in the lower panel. Unedited codons are indicated (black), while synonymous codons (upper panel only) and stop codons are highlighted with darker framed outlines.

The heatmap shown in Figure 2d (upper panel) shows relative representation of all 1,280 codon variants in the unselected library and across the dose-response profile. We observed >100 resistance-conferring edits that were enriched following drug selection (Fig. 2d). By averaging read counts for sets of synonymous codons, we derived a second heatmap showing the behaviour of all 400 possible amino acid variants across the dose-response profile (Fig. 2d, lower panel), revealing forty-six resistance-conferring edits that were enriched following drug selection; >0.1% of all normalised reads at that site following 8 nM selection.

Those edits associated with drug resistance show that oligo targeting can be used to introduce all twelve possible base edits, with no apparent bias in terms of single-nucleotide edits (SNEs), double-nucleotide edits (DNEs), or triple-nucleotide edits (TNEs). Eight of the twenty sites targeted yielded drug edits associated with drug resistance, with G^98^, Y^113^, V^128^ and S^132^ emerging as resistance ‘hotspots’ (Fig. 2d). Conversion to asparagine yielded resistance in six cases and glutamine yielded resistance in four, while only conversion to proline or arginine failed to yield resistance.

We next selected a set of nine drug-resistant mutants to interrogate the predictive power of the MOT-library profile. We used the sequencing read count data to reconstruct virtual dose-response curves, and to derive virtual EC_50_ values for these mutants (Fig. 3a). As can also be observed in Fig. 2d, codons for the same amino acid edit yielded highly consistent results. A bespoke set of ssODNs was used to introduce each selected codon-edit in otherwise wild-type cells, and a panel of mutants was assembled; a pair of independent clones for each edit. The mutants, comprising six SNEs (G^98^A, G*G*A-G*C*A; Y^113^H, *T*AC-*C*AC; D^115^N, *G*AC-*A*AC; D^116^E, GA*C*-GA*A*; V^128^M, *G*TG-*A*TG; and Y^136^F, T*A*T-T*T*T), a DNE (V^128^Q *GT*G-*CA*G) and two TNEs (Y^113^G, *TAC*-*GGA*, S^132^M *TCC*-*ATG*), were validated by sequencing, confirming the expected edits in every case (Fig. 3b; Supplementary Fig. 3); these results also indicated that heterozygous edits were sufficient to yield resistance in every case. The mutant panel was then used to derive conventional dose-response curves and EC_50_ values (Fig. 3c; Supplementary Fig. 4). A comparison with the virtual EC_50_ data revealed an excellent correlation (*R*^2^ = 0.98), indicating that codon variant scores from MOT libraries were indeed predictive of relative resistance observed using the bespoke set of mutants, and across a wide range of EC_50_ values (Fig. 3d). Thus, we identified forty-six distinct drug-resistant mutants, which displayed up to 70-fold increased resistance relative to wild-type cells, and which included all four previously known drug resistance mutations in trypanosomatid proteasome *β5* subunits; Y^113^F in *T. brucei* ^23^, D^115^N in *T. cruzi* ^25^, and G^98^C and G^98^S in *Leishmania* ^7^. We concluded that MOT-library screening can be used to identify large numbers of distinct drug resistance associated edits in trypanosomes, and also that sequence read counts for variant codons across a dose-response profile are excellent predictors of relative resistance.

**Fig. 3:**
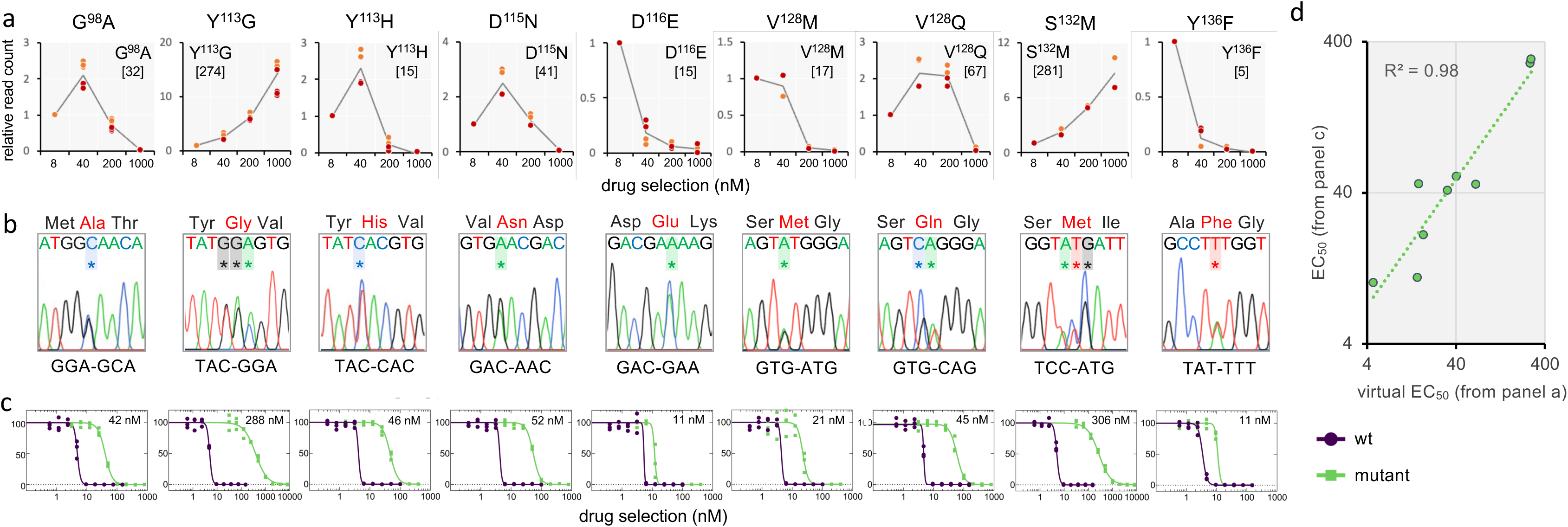
Drug resistance assays reveal the predictive power of MOT-library profiling. **a** The sequencing read counts underpinning codon variant scores were used to derive virtual dose-response curves for a selected set of edits; twenty codon edits representing nine distinct amino acid edits. We also derived a weighted average value from these data to yield virtual EC_50_ values for compound DDD247 (nM), shown in square brackets. **b** A cognate panel of mutant clones was generated using oligo targeting in a wild-type background and all clones were validated by sequencing; data for one clone is shown here, and extended data for a second set of independent clones are shown in Supplementary Fig. 3. All edited bases are indicated with an asterisk. **c** Dose-response curves and EC_50_ values for the panel of mutants. All compound DDD247 dose responses were measured in triplicate and representative dose-response curves for one biological replicate are shown here; data for the second set of biological replicates are shown in Supplementary Fig. 4. Average EC_50_ values from both biological replicates are indicated. **d** The plot compares virtual EC_50_ values from MOT-library profiling in a and average EC_50_ values from the bespoke panel of mutants in c, which are strongly correlated, *R*^2^ = 0.98.

As a further test of MOT-library profiling, we screened the libraries used above with bortezomib, which binds a site immediately adjacent to the DDD247 binding site (Supplementary Fig. 5a). Bortezomib is a modified Phe-Leu dipeptide with a boronic acid warhead ^23^; the compound is approved for cancer chemotherapy, but resistance is a significant problem ^21^. Selection of the MOT-library with 4 nM bortezomib, approximately twice the EC_50_ for wild-type *T. brucei*, yielded resistant cells, while selection with higher drug concentrations failed to do so. Amplicon sequencing and codon variant scoring revealed only two conservative amino acid edits associated with bortezomib resistance, M^97^I and M^97^V (Supplementary Fig. 5b); edits that notably failed to yield resistance to compound DDD247. These results highlight the ability of MOT-library screens to discriminate between inhibitors that engage immediately adjacent sites within the same target.

### Constraints imposed on mutational resistance space by fitness space

More than 90 % of edits fail to yield drug resistance in our screens, either because they fail to impact drug binding affinity, or simply because they yield defective proteasomes. Thus, mutational resistance space will be constrained by mutational fitness space at any given drug target binding site. In order to assess mutational fitness space at the proteasome β5 subunit drug binding site, we deleted one copy of the *β5* gene, assembled MOT-libraries in the diploid and haploid strains, and derived codon variant scores after six hours and four days, without any drug selection (Fig. 4a). To validate the approach, we first assessed edits producing stop codons, which were severely depleted in the haploid strain, but could be retained in the diploid strain, as expected (Fig. 4b). This analysis also revealed dominant-negative impacts of more distal stop codons in the diploid strain, likely due to the expression of longer, albeit truncated, β5 peptides.

**Fig. 4:**
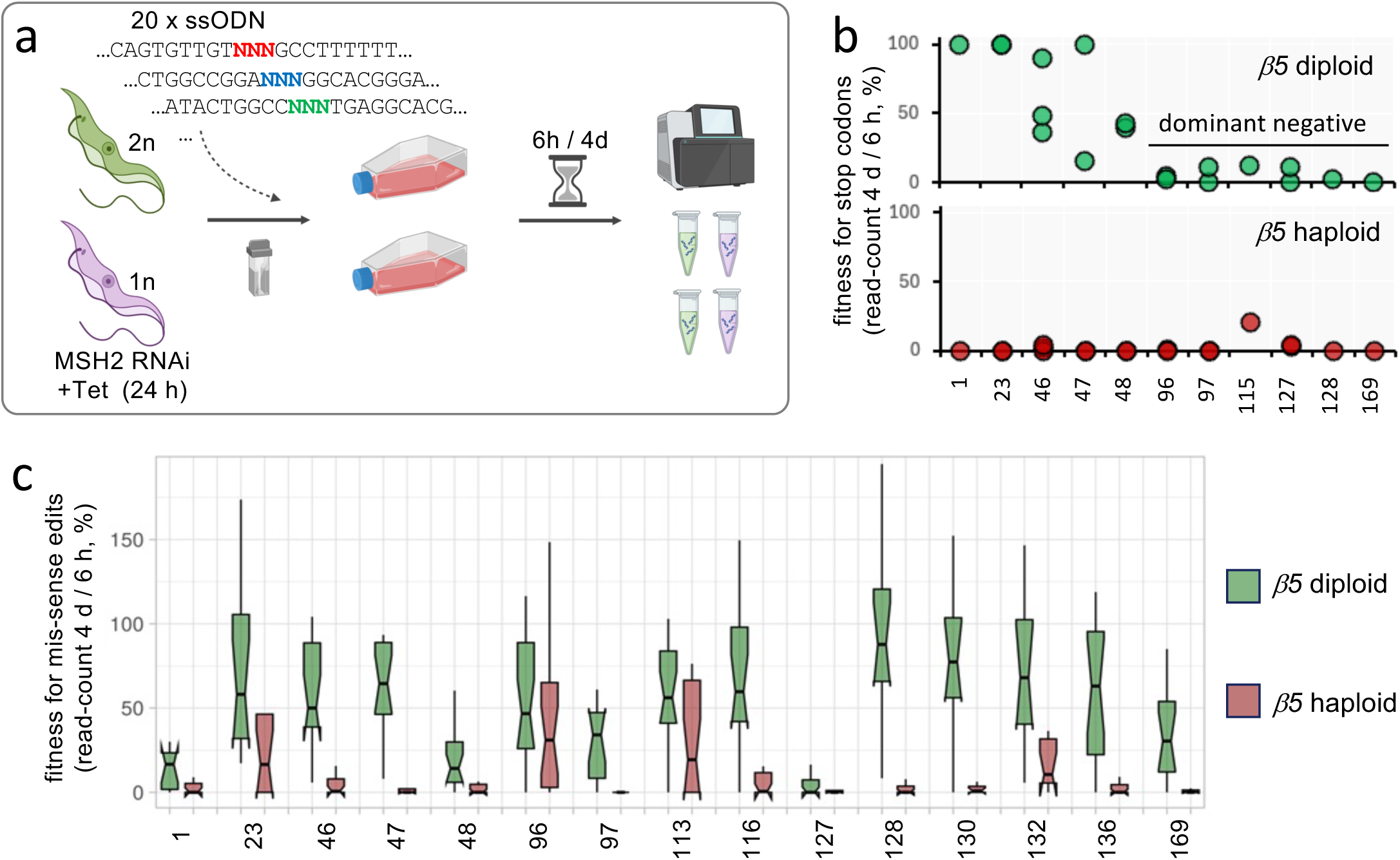
Constraints imposed on mutational resistance space by fitness space. **a** The schematic illustrates the MOT-library fitness profiling approach. The set of twenty ssODN with degenerate codons were individually transfected into *T. brucei MSH2* RNAi cells, either with two (diploid, 2n, green) or one (haploid, 1n, purple) proteasome *β5* allele, 24 h after inducing knockdown with tetracycline. Cultures were assessed using deep sequencing of *β5* amplicons and codon variant scoring prior to transfection, 6 h after transfection, and in duplicate 4 d after transfection, for each genotype. Created in BioRender. Altmann, S. (2025) https://BioRender.com/wi23cs9 **b** The plots show fitness, as revealed by the relative percentage of reads for stop codon edits remaining 4 d after ssODN transfection, and at the sites indicated in the proteasome β5 subunit; in the diploid strain (upper panel) or in the haploid strain (lower panel). **c** The boxplot shows fitness, as revealed by the relative percentage of reads for mis-sense (non-synonymous) edits remaining 4 d after ssODN transfection, at the sites indicated in the proteasome β5 subunit. Boxes indicate the interquartile range (IQR), the whiskers show the range of values within 1.5 × IQR, and a horizontal line indicates the median. The notches represent the 95% confidence interval for each median.

We next turned our attention to edits generating mis-sense (non-synonymous) codons, comparing relative codon-variant scores from the diploid and haploid strains, and for fifteen sites for which we obtained sufficient data. Mutational tolerance was reduced at all fifteen of these sites in the haploid strain relative to the diploid strain, indicating widespread loss-of-fitness associated with mutant proteasomes (Fig. 4c). Beyond this feature, three notable categories of edited codons emerged. First were codons for which most edits were poorly tolerated in both strains, consistent with dominant-negative defects (T^1^, G^48^, S^127^); second were codons for which many edits were tolerated only in the diploid strain, consistent with loss-of-proteasome function (A^46^, G^47^, M^97^, D^116^, V^128^, S^130^, Y^136^, A^169^); and third were codons for which many edits were tolerated in both strains, consistent with retained proteasome function (G^23^, S^96^, Y^113^, S^132^) (Fig. 4c). Dominant-negative edits were not expected to yield drug resistance, and indeed we saw no evidence for resistance-associated edits at these three sites, which included the catalytic threonine at T^1^ (Fig. 2d). The second loss-of-function category edits were also not expected to yield robust drug resistance, and indeed resistance-associated edits were limited at these eight sites, typically with failure to tolerate >40 nM drug selection (Fig. 2d); V^128^ did emerge as a resistance hotspot above but the mean virtual EC_50_ for edits at this site was only 23 nM (n=8). In contrast, edits in mutational fitness space, those associated with retained proteasome function, have the potential to yield robust resistance, and indeed we identified both Y^113^ and S^132^ as resistance hotspots above (Fig. 2d). Thus, although only a minority of edits around the proteasome β5 subunit drug binding site sustain fitness, a substantial proportion of these edits yield robust drug resistance. We concluded that MOT-library profiling can be employed to elucidate both drug resistance space at a given drug binding site, and constraints imposed on resistance space by fitness space.

### Accurate prediction of drug resistance mutations in SNP-accessible space

The codon-degenerate ssODNs used above were able to introduce the full range of SNEs, DNEs, and TNEs, and to access all amino acid variants. The vast majority of naturally occurring mutations are limited to single nucleotides, however, and can access only nine alternative codons, or five to seven alternative amino acids (see Fig. 5a, lower left segment). Using the MOT-library profiling data to predict naturally occurring mutations that confer compound DDD247 resistance, we found that only fifteen of forty-six amino acid edits linked to drug resistance above were accessible via an SNE (Fig. 5a, upper right segment, green boxes); six are at the Y^113^ resistance ‘hotspot’, while only DNEs and TNEs were linked to resistance at the S^132^ ‘hotspot’ (Fig. 5b).

**Fig. 5:**
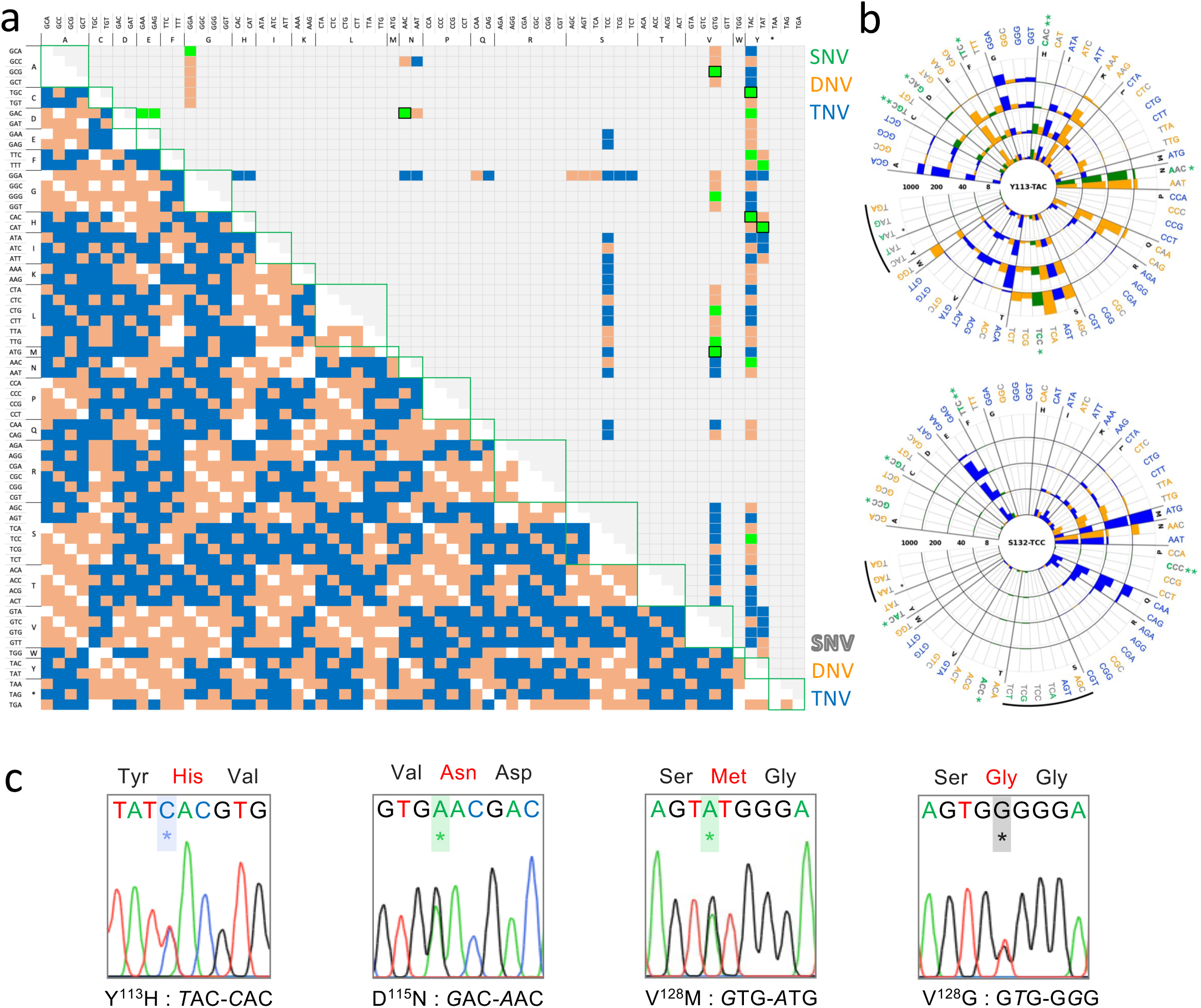
Accurate prediction of drug resistance mutations in SNP-accessible space. **a** The lower left-hand half of the plot illustrates the mutability of a given codon into any other codon via exchange of one (single nucleotide variant, SNV, white), two (double nucleotide variant, DNV, orange) or three (triple nucleotide variant, TNV, blue) nucleotides. The upper right-hand half of the plot shows those codon edits observed in our DDD247-resistance MOT screening profile, with SNVs shown in green, and transitional SNVs also framed. **b** The radial plots show dose-response data for two DDD247-resistance hotspots observed in the proteasome β5 subunit in our MOT-profiling screens (Fig. 2d), indicating that resistance is only accessible via SNVs at the Y^113^ hotspot. TAC^113Y^ can access TGC^C^, GAC^D^, TTC^F^, CAC^H^, AAC^N^ and TCC^S^ via an SNV, all predicted to yield resistance. TCC^132S^ can access GCC^A^, TGC^C^, TTC^F^, CCC^P^, ACC^T^ and TAC^Y^ via SNVs, but none of these are predicted to yield resistance. Transitions, **; transversions, *, black arcs denote synonymous and stop codons. **c** Sanger sequencing data for four spontaneous and predicted SNVs in the proteasome β5 subunit associated with DDD247-resistance. *, indicates mutated bases.

Since generation of spontaneous proteasome inhibitor-resistant trypanosomatid mutants typically takes several months ^7,17,25^, we took advantage of transient knockdown of mismatch-repair ^14^ to rapidly generate a panel of resistant *T. brucei* mutants. We sub-cloned the inducible mismatch-repair knockdown strain, induced knockdown for 24 h, subjected each clone to 40 nM drug pressure for 6-9 days, sub-cloned each resistant population, and sequenced the *β5* gene in a single clone from each population. Sequence analysis revealed that sixteen of nineteen of these independent clones had mutations in the *β5* gene, all of which were single nucleotide mutations (Fig. 5c), as expected. Thirteen were V^128^M (*G*TG-*A*TG) mutants, which may have arisen frequently due to the presence of a native editing template; we identified a potential A-mismatched template sequence (TGAAATTTTTAGT*A*T) on four distinct chromosomes. Notably, this mutant also registered a particularly strong resistance signal in Fig. 2d. Although there are twice as many possible transversions (A or G - C or T) as transitions (A - G or C - T; see Fig. 5a-b), transitions, like the V^128^M mutation, are more frequent. Indeed, assessment of 355,892 polymorphisms ^26^ suggested that transitions are 1.8 times more frequent than transversion in *T. brucei*. Accordingly, the other three *β5* subunit mutants comprised two transitions (Y^113^H, *T*AC-*C*AC; D^115^N, *G*AC-*A*AC) and a transversion (V^128^G, G*T*G-G*G*G) (Fig. 5c, Supplementary Fig. 6a), while the remaining three resistant clones had single nucleotide mutations in the proteasome *β4* subunit (Tb927.10.4710), all at F^24^L (Supplementary Fig. 6b), which also interacts with compound DDD247 ^7^.

The resistance-associated mutation profile observed in *T. brucei* using MOT-library profiling (Fig. 2d) was also predictive of those other spontaneous drug resistance mutations reported previously in trypanosomatid proteasome *β5* subunits; Y^113^F in *T. brucei* ^23^, D^115^N in *T. cruzi* ^25^, and G^98^C and G^98^S in *Leishmania* ^7^, all accessible via single nucleotide mutations; G^98^C (*G*GC-*T*GC) and G^98^S (*G*GC-*A*GC) are only accessible via single nucleotide mutations in *Leishmania* due to distinct codon-usage at this site; GGA in *T. brucei*. In summary, all spontaneous DDD247-resistant *T. brucei* clones had a single nucleotide mutation in a proteasome subunit, with 84 % in the *β5* subunit; all *β5* subunit mutations were predicted by MOT-library profiling (Fig. 2d, Fig. 5a); and no β5 subunit mutations were identified in residues more than 5 Å from the drug binding site (Fig. 1a). These observations, taken together, indicate that MOT-library profiling in *T. brucei* facilitates accurate prospective prediction of spontaneous drug resistance mutations in multiple trypanosomatids.

### Computational modelling of ligand affinity for mutant proteasomes

For computational modelling of ligand - proteasome binding affinity, we built a homology model of the *T. brucei* proteasome (Fig. 6a), revealing a high degree of similarity between the ligand binding sites in *Leishmania* ^7^ and *T. brucei* (Fig. 6b). Docking calculations focused on the cavity formed by residues within 5 Å of the DDD247 ligand (Supplementary Fig 7a-b) and yielded excellent correspondence with the published *Leishmania* / GSK3494245 structure (RMSD average: ∼0.35 Å) ^7^, while docking of compound DDD247 yielded a similar pose, with very good alignment to the aromatic and aliphatic rings (RMSD average: ∼0.35 Å) (Supplementary Fig 7c). Molecular dynamics simulations of the complex with compound DDD247 revealed stabilisation of ligand-protein interactions, by hydrogen bonding with β5 residues, Y^113^, G^129^, and S^130^, in particular, and with other hydrophobic interactions also present along the simulation, with β5-Y^113^ and β4-F^24^; while the morpholine group displays evidence of being rather flexible and solvent exposed (Supplementary Fig 8a-b).

**Figure 6:**
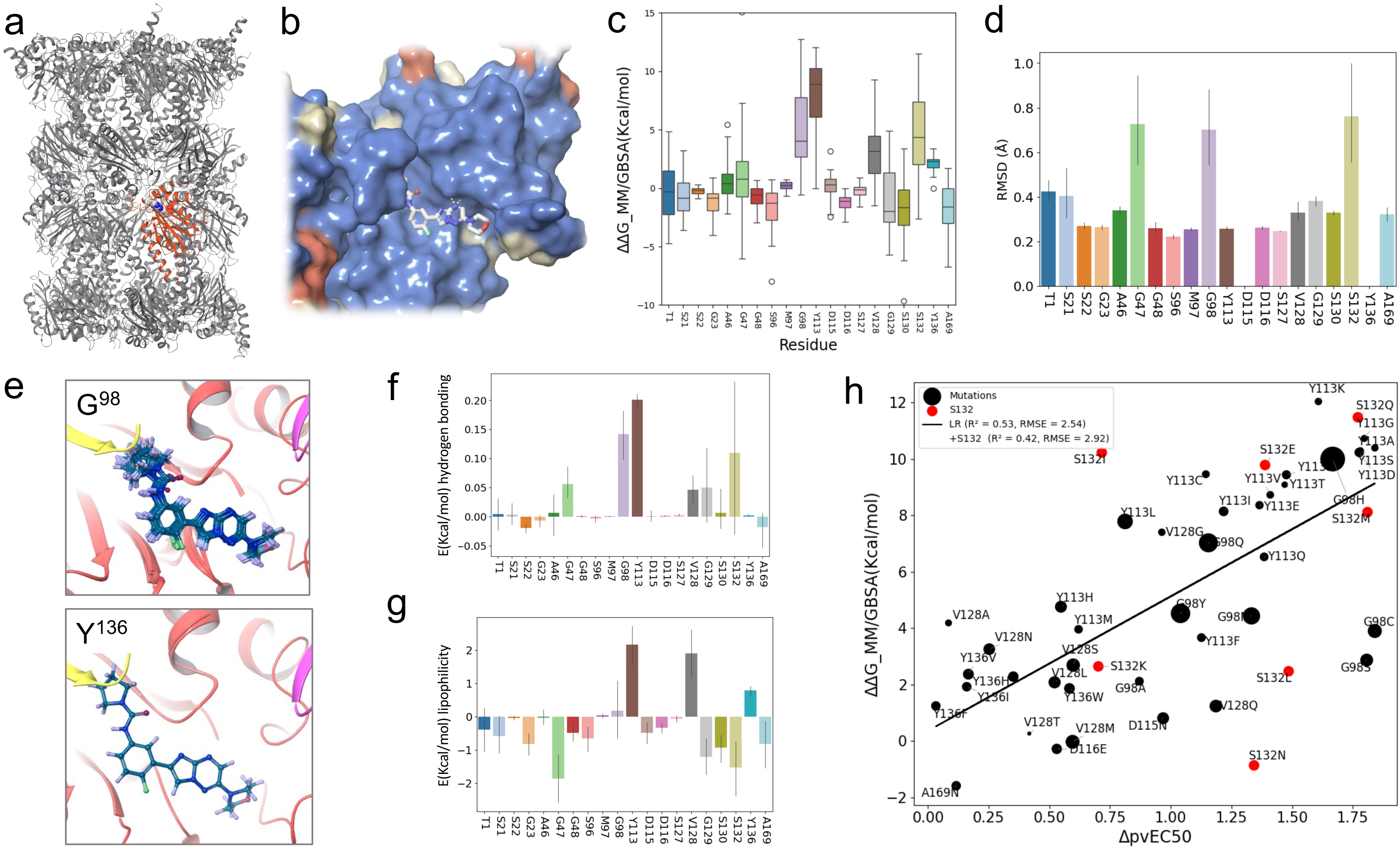
Computational modelling of ligand affinity for mutant proteasomes. **a** The *T. brucei* proteasome homology model, highlighting the β5 subunit in red and the DDD247 ligand binding site. **b** The ligand DDD247 binding site is surrounded by residues that are conserved between *Leishmania* and *T. brucei*. Conserved, blue; similar, yellow; dissimilar, orange. **c** The boxplot shows the impact of mutations on ligand affinity at the sites indicated, as predicted using MM/GSA free energy calculations. Boxes indicate the interquartile range (IQR), the whiskers show the range of values within 1.5 × IQR, and a horizontal line indicates the mean. **d** RMSD for the ligand predicted for mutations at each site. Values indicate the mean with SD for all mutations relative to the native conformation. **e** The examples show the impact of mutations at G^98^ and Y^136^, where the ligand is overlaid from the final minimisation of all possible mutations. **f** The calculated binding free energy from c was decomposed into hydrogen bond contributions. Values indicate the mean with SD for all mutations relative to the native conformation. **g** As in f but for lipophilicity. **h** Correlation between binding affinity (ΔG_MM/GBSA) and virtual EC_50_ values calculated from MOT-profiling data (ΔpvEC_50_). *R*^2^ is 0.42 for all sixty-four mutations analysed, and 0.53 when S^132^ mutations are excluded (red). Datapoint size indicates SD for clusters in set 1, while data for S^132^ mutations were derived from cluster 8 (Supplementary Fig 9).

We next considered all possible mutations in those twenty β5 subunit residues within 5 Å of the DDD247 ligand. After clustering snapshots from the molecular dynamics simulation trajectory (Supplementary Fig 9), selected clusters were used for ΔΔG Molecular Mechanics General Born Surface Area (MM/GBSA) calculations ^27^, revealing G^98^, Y^113^, V^128^, and S^132^ as hotspots where mutations were predicted to decrease ligand affinity (Fig. 6c). Remarkably, these were the same four hotspots identified by *in cellulo* MOT-profiling above (Fig. 2d). Among these hotspots, we observe ligand displacement at G^98^ and S^132^ (Fig. 6d). Indeed, the resistance-associated mutations we observe *in cellulo* all increase the side-chain mass at these sites; by 58 +/- 31 Da at G^98^ (n = 7) and by 35 +/- 8 Da at S^132^ (n = 7). Exemplar images illustrate how G^98^ and Y^136^ mutations have substantial or minimal impacts on ligand binding, respectively (Fig. 6e).

Hydrogen-bonding (Fig. 6f) and lipophilicity (Fig. 6g) are also predicted to have substantial impacts of protein-ligand affinity. Both features are predicted to contribute to the impact of Y^113^ mutations, since this residue can form a hydrogen bond with the ligand and also interact within a lipophilic environment. Mutations at G^98^, although they do not form a hydrogen bond with the ligand, all disrupt hydrogen bonding between the adjacent Y^113^ residue and the ligand, while impacts on lipophilicity appear to be the major drivers of drug resistance for V^128^ mutations. In the case of S^132^, molecular dynamics clusters often placed this residue distal from the DDD247 ligand binding pose. Cluster 8, however, indicated a direct hydrogen bond between S^132^ and the ligand, and this bond is lost in mutations associated with drug resistance. Accordingly, we suggest that this hydrogen bond does occur but is not observed in the cryo-EM structure. By comparing these ΔΔG (MM/GBSA) calculations with the full set of forty-six virtual EC_50_ values derived above using MOT-profiling, we obtained an *R*² value of 0.42 (Fig. 6h). As an example of excellent correlation between observation *in cellulo* and prediction *in silico*, Y^113^ mutations incorporating highly charged residues (Y^113^K, Y^113^D, Y^113^Q, Y^113^S) are associated with a higher degree of resistance relative to mutations that conserve hydrophobicity (Y^113^F, Y^113^L, Y^113^H, Y^113^M).

### Models that combine affinity and function parameters improve resistance predictions

We elucidate above the mutational fitness space that restricts mutational resistance space at the DDD247 binding site; whereby non-functional mutant proteasomes fail to yield drug-resistant cells (Fig. 4). We, therefore, sought to combine predictions of loss of protein function with our ligand affinity predictions. We used Evo 2, a DNA language model trained on >9 trillion DNA base pairs from all domains of life ^28^, to predict loss of proteasome function due to codon changes, considering the full set of 1,280 mutations surveyed by MOT-library profiling above. Evo2 Δ_scores for stop codons were used to define a threshold, below which a mutated β5 subunit was considered non-functional (Supplementary Fig 10a) and scores were then plotted for mutations at each site (Supplementary Fig 10b). Both the affinity predictions (Fig. 7a, left-hand panel) and functional predictions (Fig. 7a, right-hand panel) were then combined to generate a heatmap showing resistance predictions (Fig. 7a, lower panel). The resistance hotspots described above, G^98^, Y^113^, V^128^ and S^132^, were retained, and the predicted impact of each individual mutation can now be seen in the combined heatmap. Although the analysis predicted reduced ligand affinity due to mutations at A^46^, G^47^, and G^129^, loss of function is also predicted due to these mutations, meaning that resistance is predicted to be minimal (Fig. 7a). Consistent with these predictions, no mutations at these sites registered as resistance mutations in our *in cellulo* screen (Fig. 2d); a circular plot shows the full *in cellulo* resistance profile and *in silico* predicted resistance profile side-by-side for comparison (Fig. 7b). These results highlight the potential for added value when combining affinity and function parameters to predict drug resistance.

**Figure 7:**
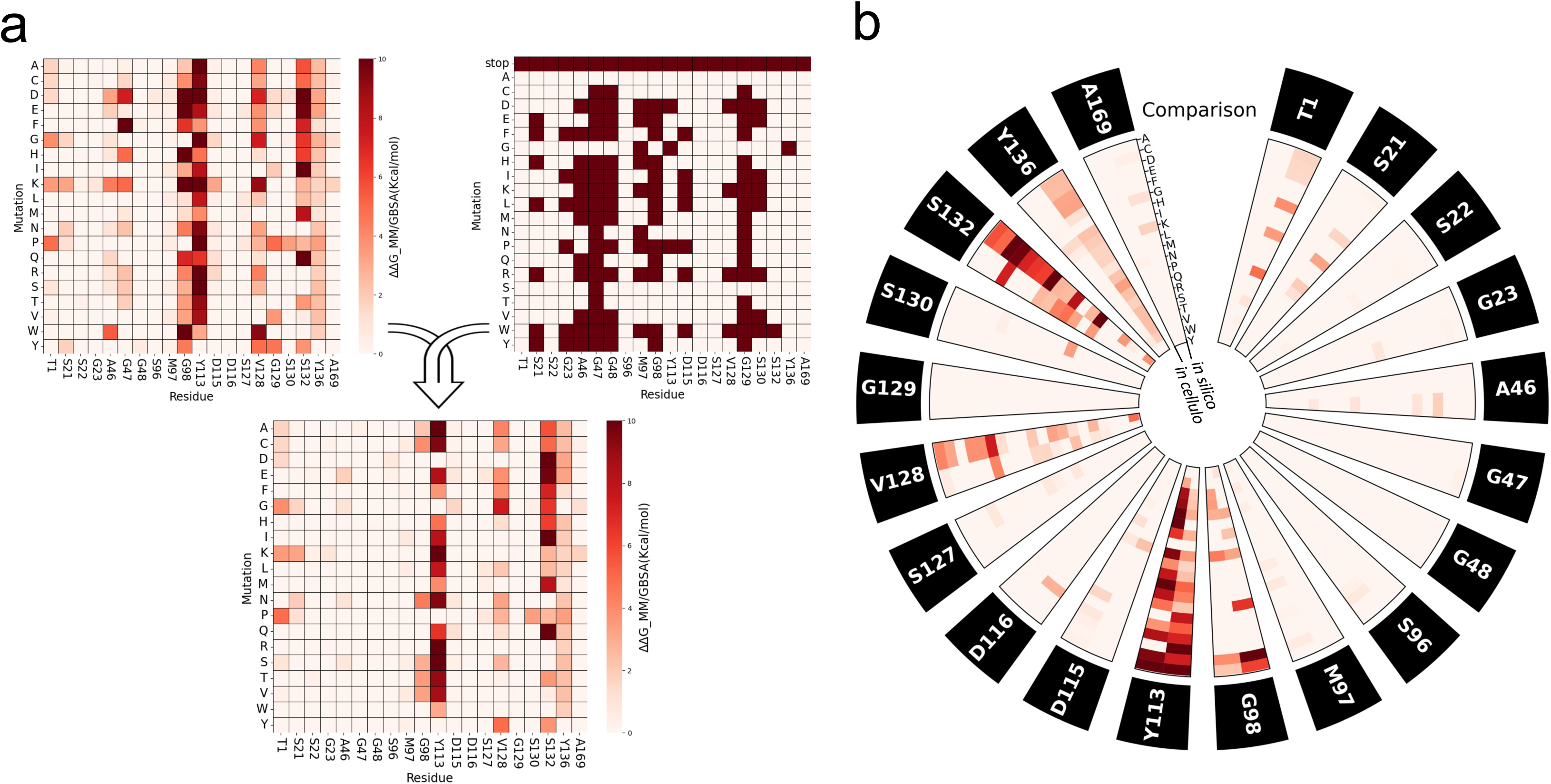
Computational modelling combining affinity and function parameters. **a** The top left heatmap shows the impact of the full set of mutations on ligand affinity, as predicted using minimisation and MM/GBSA calculations, normalised to 0 as the baseline for the native sequence. The top right map shows the impact of the full set of mutations on proteasome function, as predicted using Evo 2; dark shade indicates loss-of-function. The lower heatmap shows predicted DDD247-resistance as determined by combining loss of ligand affinity predictions with loss of protein function predictions. Mutations with predicted loss of function were set to 0. **b** The circular plot compares the *in cellulo* measures of DDD247-resistance (virtual EC_50_ values) from Fig. 3a and the *in silico* predicted measures of DDD247-resistance from the lower heatmap in a.

## Discussion

Key goals in drug discovery are to evolve potent, safe, and durable therapies. The optimisation of structure-activity relationships typically requires substantial investment, however, with medicinal chemistry often informed by structural data and modelling when the target is known. Despite the importance of understanding structure-activity relationships, and their impacts on efficacy and resistance, the relevant interactions often remain hypothetical, speculative, and incompletely characterised; in particular, empirical in *cellulo* data regarding individual residues at a drug binding site are often lacking.

To address this knowledge gap, we sought to scale the oligo targeting approach in *T. brucei* ^14^. We focused on the proteasome, an established drug target in diverse cell types ^7,17,20–22^, and developed Multiplex Oligo Targeting for site saturation mutagenesis. Guided by cryo-electron microscopy structures of the ligand-bound proteasome ^7^, we identified residues within 5Å of the ligand, and constructed libraries of cells with all possible mutations at these sites. Following drug selection, we profiled all drug resistance edits using deep sequencing and variant scoring. The genome editing approach used here is rapid and precise, allows for expression of mutants from the otherwise unperturbed native locus, and offers high fidelity multiplexed assessment of edits quantified directly at the target locus. We also note that the heterozygous edits we observe most likely reflect those potential routes to drug resistance in a clinical setting. Thus, the approach facilitates decoding of efficacy and resistance space at a given drug binding site.

Since drug resistance space is constrained by fitness space, we also assessed the fitness of cells exclusively expressing edited proteasomes. The majority of proteasome edits came with a fitness cost, and indeed, these edits failed to yield robust resistance. Since spontaneous drug resistance space is further constrained by accessible mutational space, a better understanding of these parameters will facilitate prediction and surveillance, as well as the prospective design of second-line therapies that bypass anticipated resistance mechanisms. Ultimately, and highlighting the potential value of such insights, compounds that target sites within highly restricted fitness space and otherwise inaccessible resistance space should deliver more durable therapies. Compounds identified as particularly vulnerable to spontaneous resistance could also be prioritised for use in combination therapies ^29^.

Computational models and artificial intelligence-based approaches promise to revolutionise the design of more active and durable drugs. *In silico* predictions often remain unvalidated due to the limited availability of *in cellulo* experimental data, however. By combining empirical data with *in silico* predictions of drug target affinity and proteasome structure - function perturbing mutations, we demonstrated how effective *in silico* predictions can be. Notwithstanding some limitations, the computational models we present yield effective *in silico* predictions of drug target affinity and relative resistance, explained by a combination of steric effects, hydrogen-bonding and lipophilicity. Such models will require comprehensive, high-quality training data to further improve their utility in future drug design.

Bridging the knowledge gap in relation to mutations associated with drug efficacy and resistance is a priority, especially given rapid advances in the complementary areas of structural biology and computational drug design. Specifically, several new anti-microbial drugs with known targets are now in pre-clinical or clinical development ^30,31^, and we aim to improve our understanding in this area. By assessing and understanding the impacts of all possible mutations at a range of drug binding sites, we can improve our understanding of mutations associated with drug resistance, improve prospects for resistance surveillance and disease treatment, and enhance future computational drug design strategies.

## Methods

### *T. brucei* growth and manipulation

Bloodstream form Lister 427 *Trypanosoma brucei* wild-type and *MSH2* RNAi ^14^ strains were cultured in HMI-11 (Gibco) supplemented with 10 % fetal bovine serum (Sigma) at 37°C and with 5 % CO_2_ in a humidified incubator. Genetic manipulations was carried out by electroporation using a Nucleofector (Lonza), with a Human T-cell kit (Lonza), or with custom-made transfection buffer ^32^, with the Nucleofector set to Z-001 (Amaxa).

### Oligo targeting

Oligo targeting was carried out using single-stranded oligodeoxynucleotides (ssODNs, Thermo Fisher Scientific) essentially as described ^14^. Briefly, for routine assays, we used 40 μg of each ssODNs in 10 ul of 10 mM Tris-Cl, pH 8.5, mixed with 25 million cells in 100 μl transfection buffer. Selection with DDD247 or DDD248, compounds 7a and 7, respectively in Wyllie *et al* ^7^, was applied 6 h later, as appropriate, and cells were subjected to limiting dilution in 96-well plates as required. Genomic DNA was extracted using a Qiagen DNeasy Kit followed by PCR amplification of the β5 gene using KOD Hot Start DNA Polymerase (Merck) or Phusion polymerase (NEB). PCR products were purified using a PCR Purification Kit (Qiagen) and, to identify edits, amplicons were subjected to either Sanger sequencing (Genewiz, Azenta Life Sciences) or deep sequencing using DNBseq (BGI Genomics).

### Assembly and screening of multiplexed oligo targeting libraries

For site saturation mutagenesis using a set of twenty degenerate ssODNs, we used 40 μg of each ssODNs in 10 ul of 10 mM Tris-Cl, pH 8.5, mixed with 6.5 million cells in 100 μl transfection buffer. *MSH2* knockdown with induced 24 h prior to transfection by the addition of 1 μg/mL tetracycline (Sigma). Each ssODN was transfected individually and the cultures were then pooled in 150 ml of medium to generate the library. After 6 h, cells from 20 ml of the culture were cryo-preserved, cells from another 20 ml were collected for (pre-selection) DNA extraction, and the remainder were split into two cultures and subjected to selection with 8 nM compound DDD247. Following 4-5 d, 50 ml of each culture was collected for DNA extraction and the remainder was split into three cultures that were subjected to selection with either 40 nM, 200 nM or 1 uM compound DDD247. Following 2-3 d, each culture was collected for DNA extraction.

For selection with bortezomib (Sigma), a cryo-preserved library sample was thawed, allowed to recover for 24 h, and split into two parallel cultures. Bortezomib selection was applied at 4 nM (EC_50_ = 2.2 nM). Following 7-9 d, cultures were collected for DNA extraction and the remainder was split into three cultures that were subjected to selection with higher concentrations of bortezomib; none of these latter cultures yielded resistant cells.

For fitness profiling, we first generated a *T. brucei MSH2* RNAi strain with a single β5 allele. A construct was synthesized comprising a *BLA* (blasticidin deaminase) selectable marker, flanked by *T. brucei* tubulin mRNA processing sequences and ∼300 bp of *β5* gene flanking sequence to promote homologous recombination. This cassette was excised from the plasmid using AscI and used to transfect *MSH2* RNAi cells. *BLA* selection was applied at 10 μg/ml after 6 h and positive clones were assessed using PCR assays. Multiplex oligo targeting libraries were then assembled in both strains, either haploid or diploid for the *β5* gene, as described above, and again using the same set of twenty degenerate ssODNs.

### Amplicon sequencing and codon variant scoring

Deep sequencing of β5 gene amplicons was carried out using DNBseq (BGI Genomics). Filtering of sequencing reads was performed using the SOAPnuke software with parameters: “-n 0.001 -l 10 -q 0.4 --adaMR 0.25 --ada_trim --minReadLen150” for drug selection experiments and “-n 0.01 -l 20 -q 0.3 --adaMR 0.25 --ada_trim --polyX 50 -- minReadLen 150” for the haploid v diploid experiment.

Codon variant scoring was performed with the OligoSeeker (0.0.5) Python package ^33^ designed to process paired FASTQ files and count occurrences of specific codons. To visualise codon variant scores, we performed a normalization step by dividing each codon variant score by the total reads for that position and converting the fraction of reads to a percentage. This was followed by background correction whereby we subtracted the values for the control sample. Negative values were replaced with 0, and average values for the duplicate libraries were then calculated to give the codon variant scores. Edits with Codons registering elevated read-counts in the control sample, including all single nucleotide variants, were excluded from the analyses shown in Fig. 1c and Fig. 4c. To derive virtual EC_50_ values, we derived a weighted average of codon variant scores across the DDD247 compound dose range. To visualize relative codon frequency using the pyCirclize python package (https://github.com/moshi4/pyCirclize, v1.9.1), raw codon counts were normalized by the maximum values and replicates were then averaged.

### Assembly of a panel of spontaneous drug-resistant mutants

We exploited the *MSH2* RNAi strain to rapidly generate a panel of spontaneous and independent DDD247-resistant mutants. To ensure that each mutant was independent, we first used limiting dilution to isolate twenty clones from the *MSH2* RNAi cell line. *MSH2* knockdown was induced in each clone for 24 h, tetracycline was removed by washing in HMI-11 medium, and compound DDD247 was added at 40 nM. We observed cell death followed by recovery and obtained nineteen independent DDD247-resistant cultures using this approach. We sub-cloned each culture by limiting dilution, and isolated one clone from each for DNA extraction, PCR amplification of the β5 gene and Sanger sequencing (Genewiz, Azenta Life Sciences). We also subsequently PCR amplified and sequenced the proteasome β4 gene in three clones for which no mutations were identified in the β5 gene.

### Dose-response assays

To determine the Effective Concentration of drug to inhibit growth by 50 % (EC_50_), cells were plated in 96-well plates at 1 x 10^3^ cells/ml in a 2-fold serial dilution of selective drug. Plates were incubated at 37°C for 72 h, 20 ml resazurin sodium salt (AlamarBlue, Sigma) at 0.49 mM in PBS was then added to each well, and plates were incubated for a further 6 h. Fluorescence was determined using an Infinite 200 pro plate reader (Tecan) at an excitation wavelength of 540 nm and an emission wavelength of 590 nm. EC_50_ values were derived using Prism (GraphPad).

### Homology modelling, docking and molecular dynamics

A *T. brucei* proteasome homology model was generated using a cryo-EM structure of the *Leishmania* proteasome in complex with the GSK3494245 inhibitor (PDB: 6qm7) ^27^, and the homology modelling tool implemented in Maestro (Schrodinger inc.) ^34^. Docking was performed with Glide XP ^35^, using GSK3494245 for validation, followed by compound DDD247, with RMSD comparisons calculated using an MCS protocol. The complex was placed in a Cubic box with waters and ions (0.15 M) using Maestro and the system was subjected to 80 ns of molecular dynamics with Desmond ^36^. The trajectory was clustered by RMSD (resulting in 9 clusters), and representative frames were selected for further analysis. Those cluster were classified with interaction fingerprints ^37^.

### Free energy estimations

The ensemble of conformations was generated from the clusters produced by the simulation to estimate the binding free energy. The residue scanning tool implemented in Maestro was used to compute approximate binding energy scores between the ligand and wild-type/mutant proteins ^36^. This tool implements a high-throughput workflow for testing mutations with the ΔΔG associated (calculated with the MM/GBSA method) ^27^ using wild-type as reference (baseline). MM/GBSA is a free energy method that gives an approximation of relative binding free energy between states (such as ligand bound vs unbound). The method decomposes the calculation into terms; Van der Waals, Coulombic, hydrogen bonding, solvation and hydrophobic, for contributions to the final ΔG. All residues tested experimentally were mutated to build single mutants, with minimisation around 7 Å as the final step of preparation. Results were plotted with Seaborn and Matplotlib. For virtual EC_50_ values derived from MOT-profiling data, a ΔpvEC_50_ was calculated for each mutant, and plotted against the ΔΔG values.

### Protein function prediction

Machine learning approaches can effectively predict the impact of mutations on protein function ^38^ and we used an Evo 2 model to predict loss of proteasome function here (evo2_7b_base) ^28^. Taking native nucleotide sequence for the *β5* subunit gene, we considered all possible mutations in those codons surveyed by MOT-profiling. After the inference, the native sequence was used as a reference, and sequences with mutations were scored and subtracted from the reference score to obtain Evo2 Δ_scores. Scores for stop codons were used to define a threshold of -0.0135, below which mutated proteins were predicted to be non-functional. For the heatmap, we classified all values below the threshold as class 1, loss of function, and all values above the threshold as class 0. To combine the binding affinity predictions and functional predictions, and to derive potential drug resistance predictions, residues, class 1 loss of function mutants were set to 0 in the free energy heatmap. Values were then normalised to allow comparison of *in cellulo* and *in silico* predictions and the results were visualised in a circular plot with Seaborn and Matplotlib.

## Acknowledgements

We thank David Robinson for assistance with structural analysis and Anna Creelman and Hayley Bell for assistance with PCR and sequencing. This work was supported by a Wellcome Centre Award (223608/Z/21/Z) and a Wellcome Investigator Award to D.H. (217105/Z/19/Z).

## Data availability

High-throughput sequencing data generated for this study have been deposited at the Sequence Read Archive under primary accession number PRJNA1295514 (https://www.ncbi.nlm.nih.gov/bioproject/1295514).

---

**Supplementary Fig 1:**
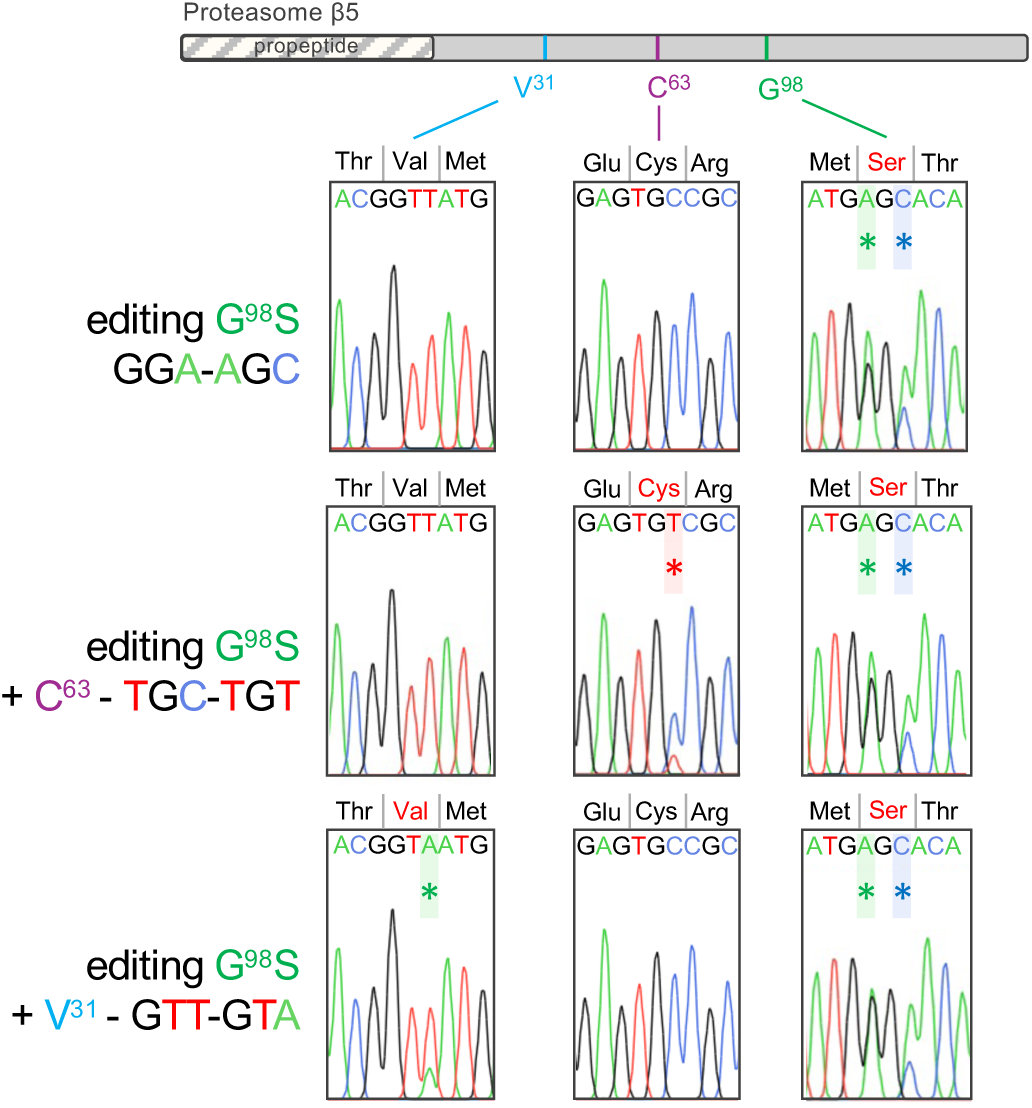
Co-editing at adjacent sites following delivery of multiple ssODNs. The schematic representation of the *T. brucei β5* sequence highlights residues targeted for editing. Sequence traces show representative outcomes following single or dual ssODN delivery and drug-selection, with 10 nM DDD248. Edited nucleotides are marked by an asterisk.

**Supplementary Fig 2:**
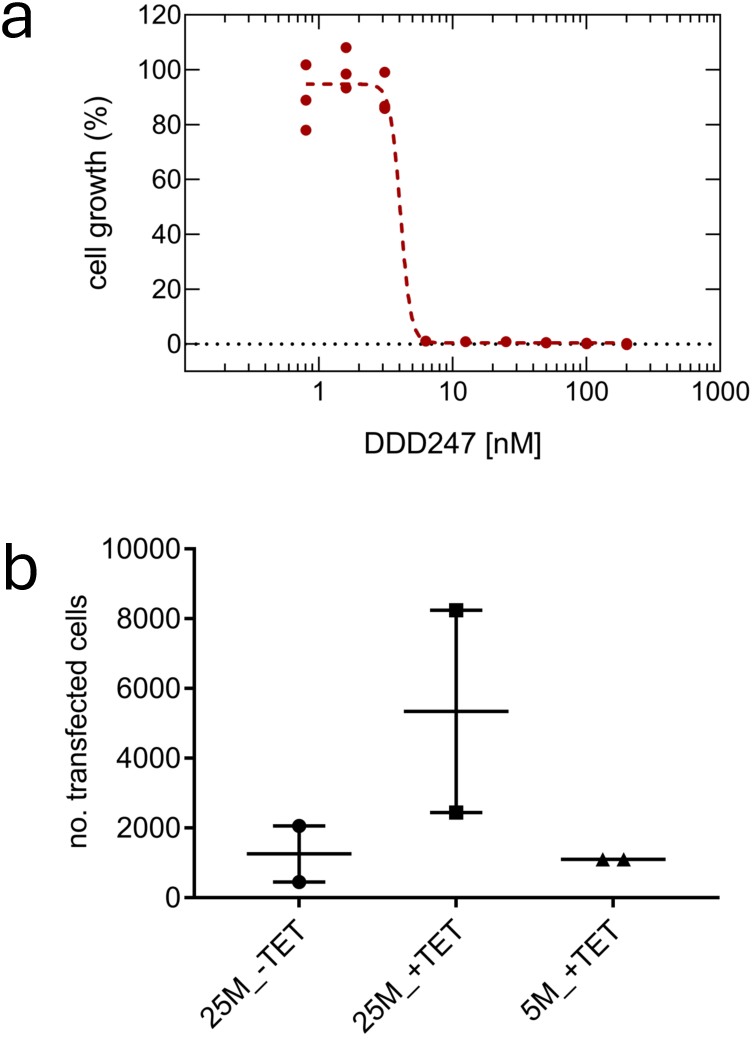
Estimation of allele replacement frequency at the *β5* gene locus. **a** The plot shows the *T. brucei* dose response curve for compound DDD247. Assays were carried out in triplicate and the EC_50_ was determined to be 4 nM. **b** An ssODN designed to introduce a GGA-AGC (G^98^S) edit and selection with compound DDD247 6 h later were used to estimate allele replacement frequency at the β5 gene locus. We tested 25 million cells with or without induction of *MSH2* knockdown 24 h prior to transfection, and also 5 million cells with induction of *MSH2* knockdown. Transfected cells were serially diluted in 96-well plates immediately after adding the drug, and no. of transfected cells was estimated 5 days later. Considering 50 % survival following transfection, the assay yielded an allele replacement frequency of approx. 0.04 % following MSH2 knockdown

**Supplementary Fig 3:**
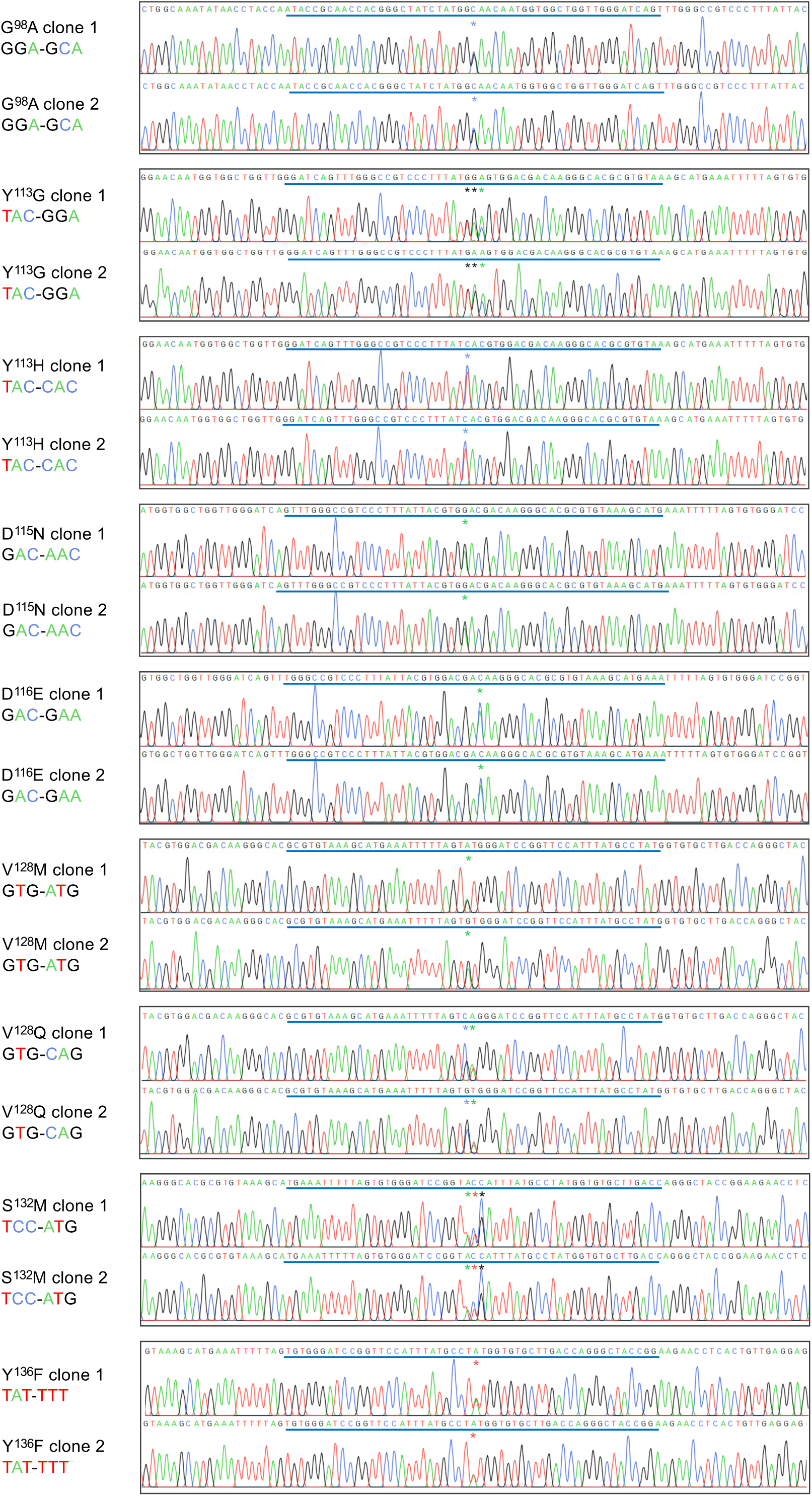
Extended sequence traces for validation of edited clones. Sanger sequence traces for nine distinct mutations introduced into wild type *T. brucei*; two independent clones of each. Edited nucleotides are marked by an asterisk. The lines above each sequence track indicates the extent of each ssODN. Other details as in Fig. 3b.

**Supplementary Fig 4:**
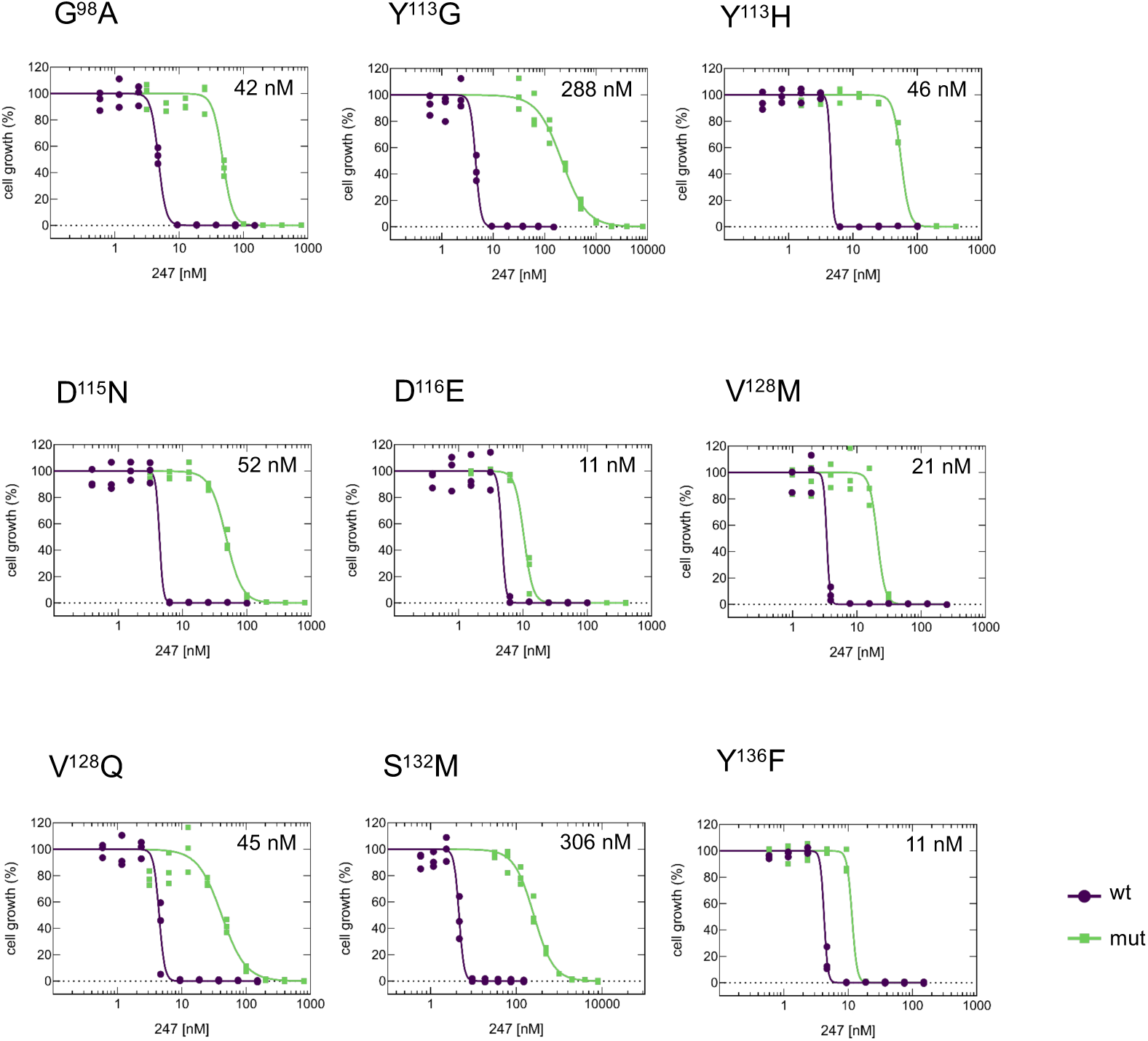
Dose response curves for edited clones. Further dose response curves for the panel of edited and drug-resistant mutants; second biological replicate for each edit. Other details as in Fig. 3c.

**Supplementary Fig 5:**
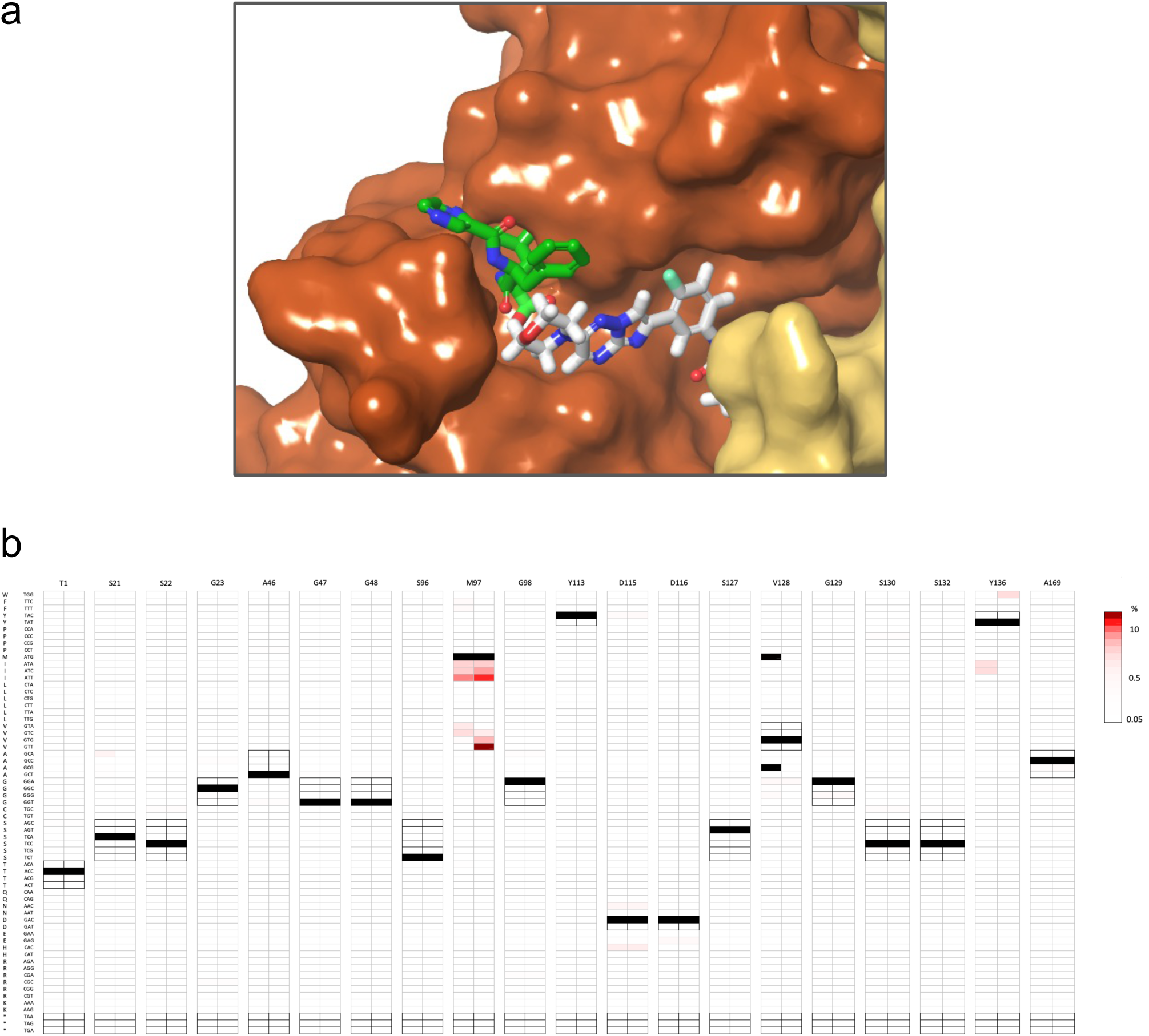
MOT-library profiling with bortezomib. **a** Docking model for DDD247 (grey) and bortezomib (green) with the *T. brucei* proteasome, indicating distinct, but adjacent binding sites. **b** Two MOT-libraries were selected with bortezomib (4nM) in parallel. All sixty-four possible codon variant scores for all twenty targeted sites in the proteasome *β5* subunit are represented as a heat-map. Unedited codons are indicated (black), while synonymous codons and stop-codons are highlighted with darker framed outlines.

**Supplementary Fig 6:**
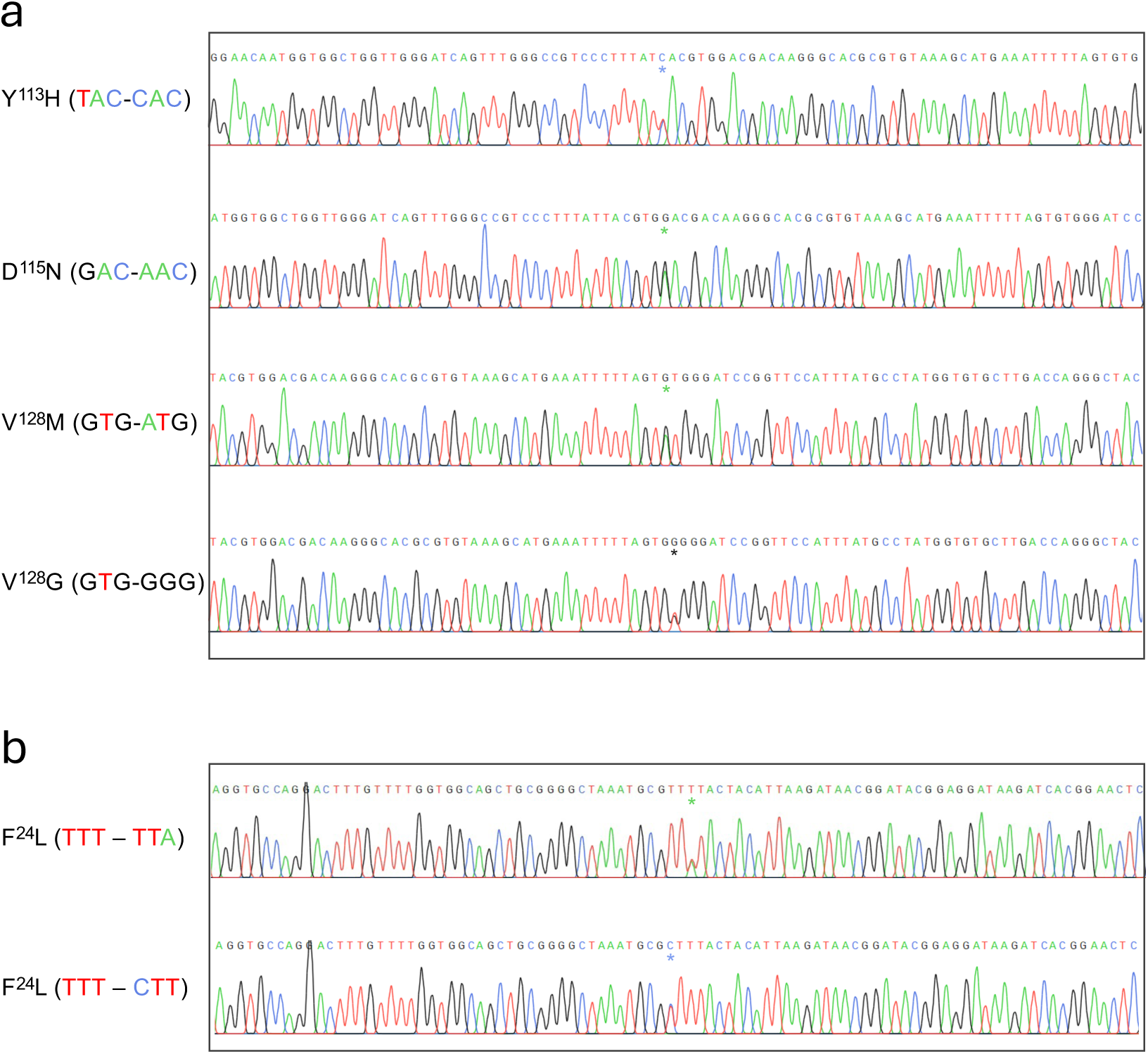
Extended sequence traces for spontaneous DDD247-resistant mutants. **a** Sanger sequencing traces for spontaneous mutations in the proteasome *β5* subunit gene following transient *MSH2* knockdown and associated with compound DDD247 resistance; all heterozygous single nucleotide variants. Mutated bases are marked by an asterisk. **b** As in a but for the proteasome *β4* subunit gene. Other details as in Fig 5c.

**Supplementary Fig. 7:**
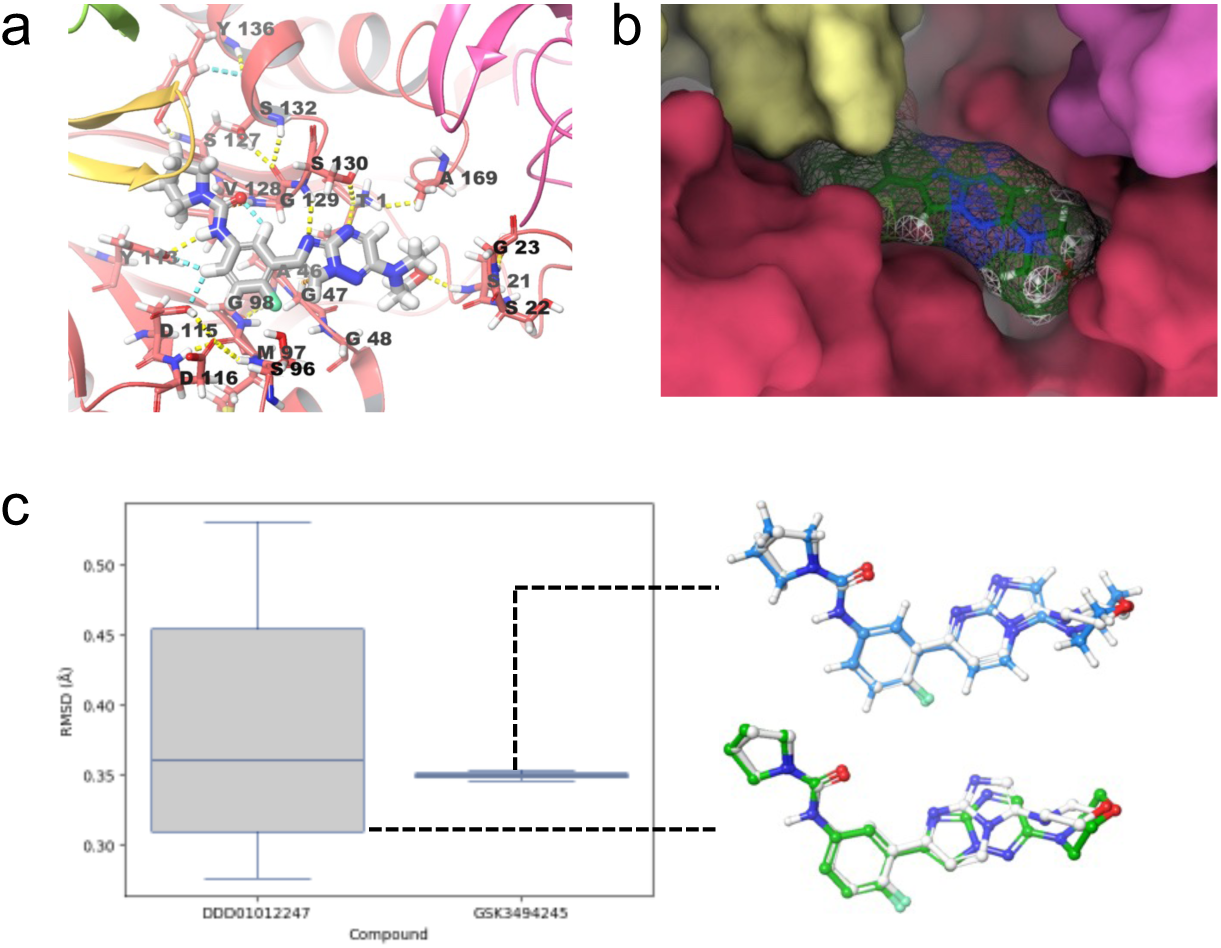
Assessment of the *T. brucei* proteasome homology model. **a** Docking model for the T. brucei proteasome and compound DDD247, indicating sites surrounded the binding pocket. **b** DDD247 sits within a well-formed cavity, primarily interacting with the β5 subunit (red). **c** RMSD values derived from comparisons of compound DDD247 (*T. brucei*) v GSK245 (pdb-Id:6qm7) and GSK245 (*T. brucei*) v GSK245 (pdb-Id:6qm7); in both cases, the reference is shown in grey.

**Supplementary Fig. 8:**
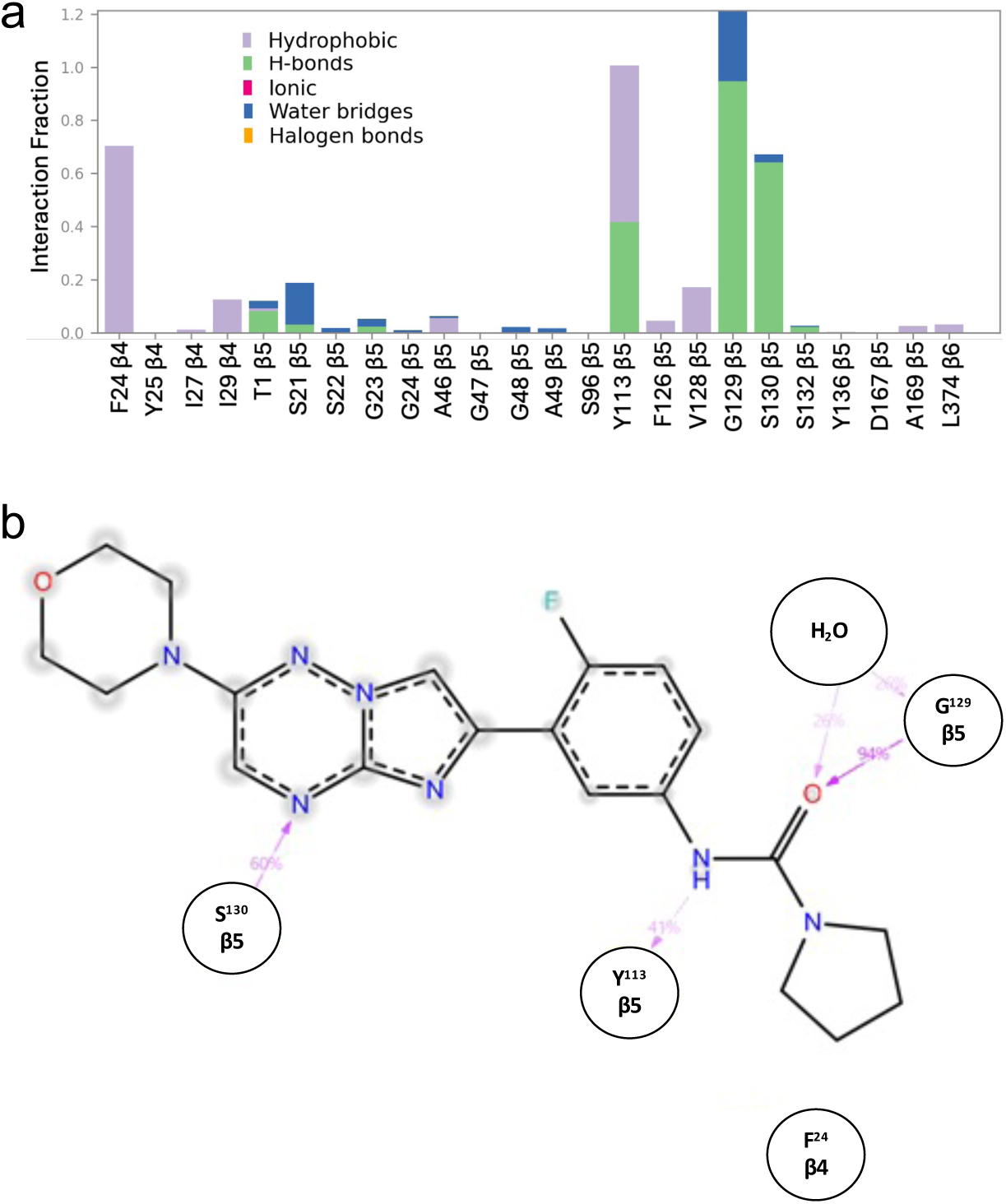
Summary of molecular interactions identified using molecular dynamics simulation. **a** The plot indicates interaction fraction and interaction class between the DDD247 ligand and subunits in the *T. brucei* proteasome homology model. **b** Percentage of major interactions along the simulation. Gray diffuse rings indicate solvent-exposed residues.

**Supplementary Fig. 9:**
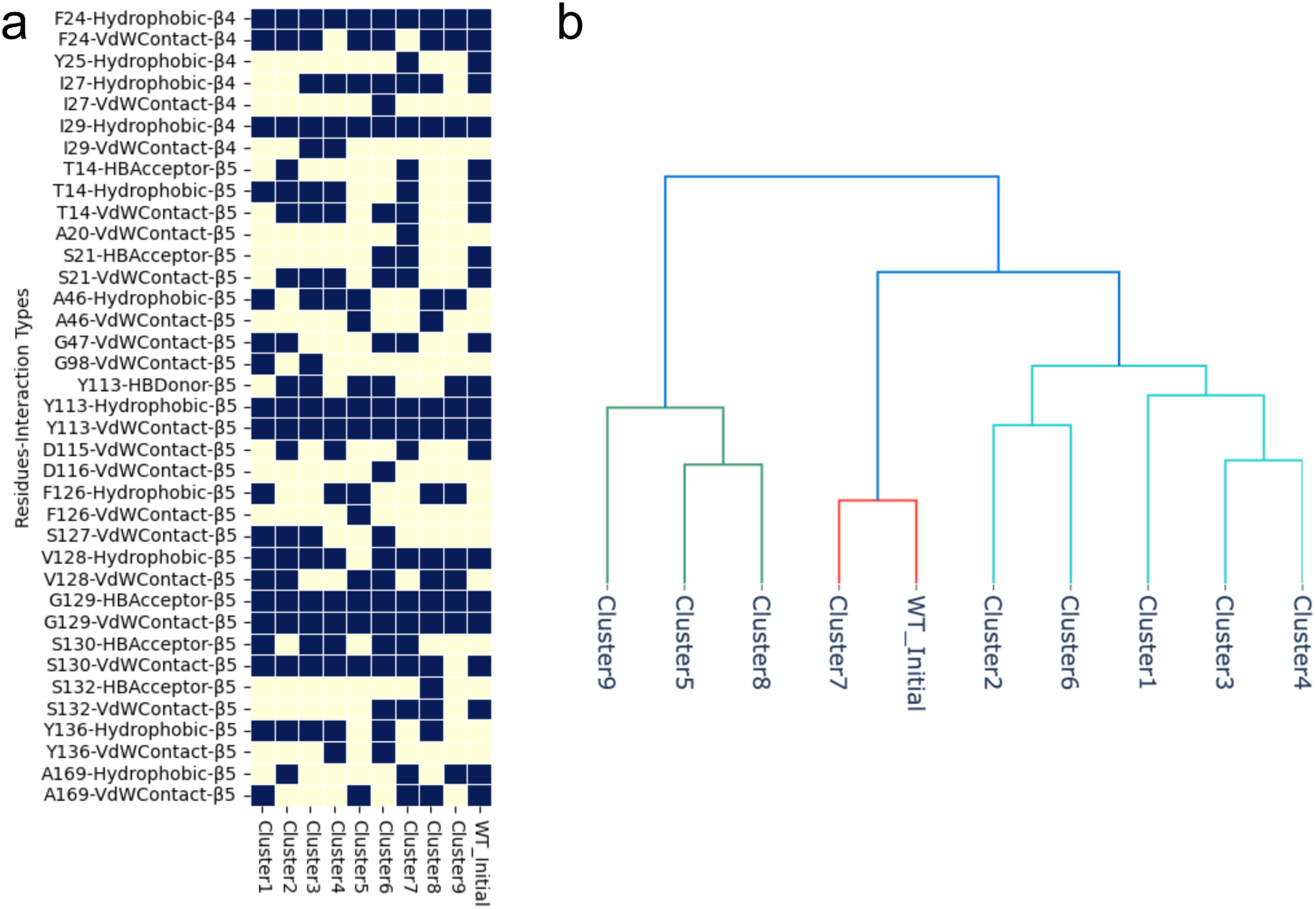
Assessment of the *T. brucei* proteasome molecular dynamics simulation. **A** Interaction fingerprints (ProLIF) of each representative snapshot from clustering; dark blue indicates interactions with DDD247. **b** Fingerprint Tanimoto similarity tree for representative snapshots from each cluster. Cluster Set 1 (2,3,6,WT_initial) and cluster 8 were selected for further analysis; cluster 8 specifically to derive data for β5 subunit S^132^ mutations.

**Supplementary Fig. 10:**
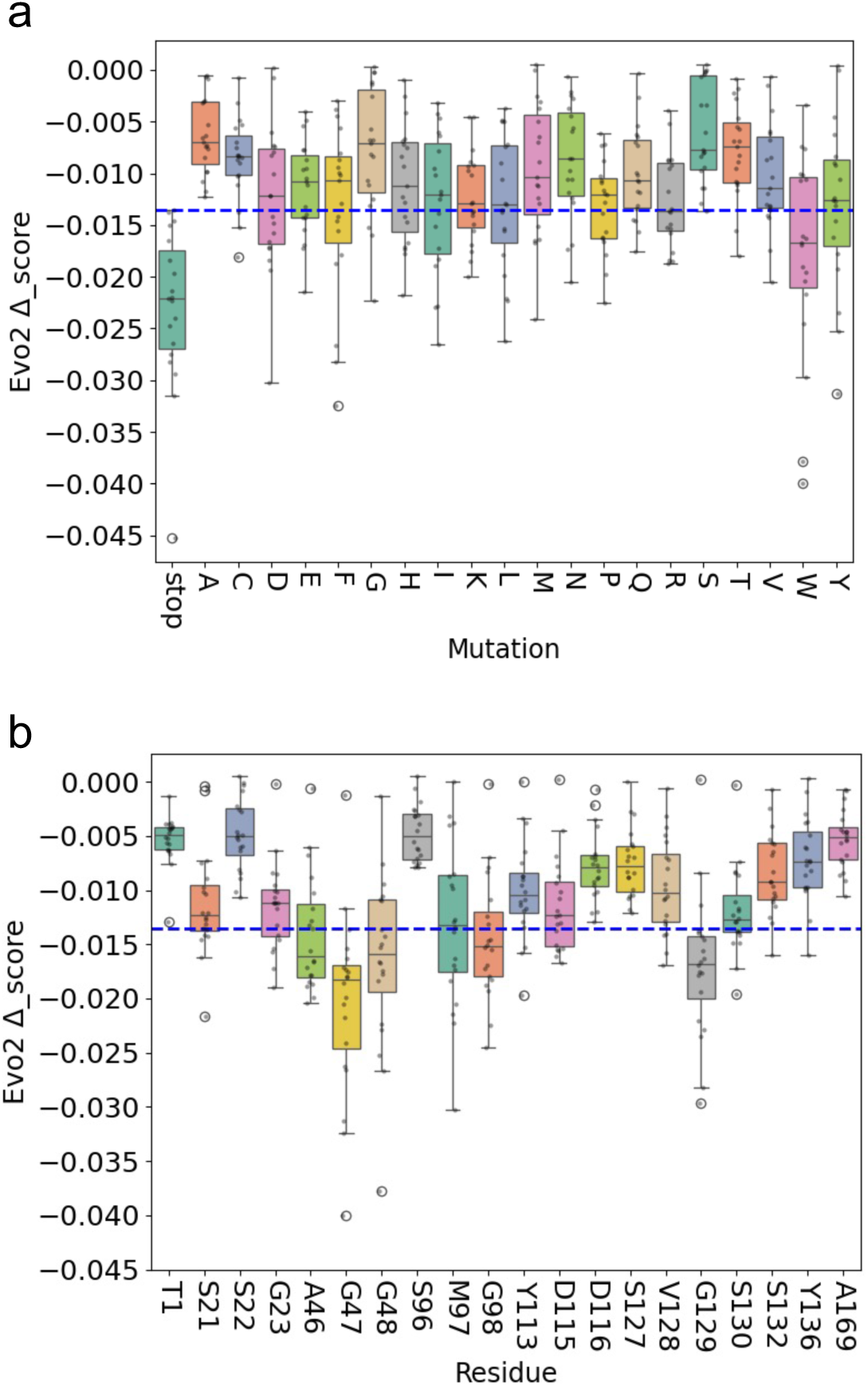
Functional predictions using Evo 2. **a** The boxplot shows Evo 2 predictions for mutations yielding each possible amino acid, across the twenty proteasome β5 subunit residues surveyed by MOT-profiling. The average Evo2 Δ_scores for codons encoding the same amino at each site are shown. The threshold for predicting function was selected based on the range of predictions for stop codons; values below this threshold, −0.0135 (blue line) were considered to indicate loss-of-function. Boxes indicate the interquartile range (IQR), and the whiskers show the range of values within 1.5×IQR. **b** The boxplot shows Evo 2 predictions for mutations yielding all possible amino acid changes at the twenty proteasome β5 subunit residues surveyed by MOT-profiling. Other details as in a.

## Notes

### Competing Interest Statement

The authors have declared no competing interest.

